# Wingless promotes JNK/MMPs positive feedback loop mediate tumour microtubes expansion, glioma progression and neurodegeneration

**DOI:** 10.1101/520346

**Authors:** Marta Portela, Natasha Fahey-Lozano, Sergio Casas-Tintó

## Abstract

Glial cells display a network of projections (cytonemes) which mediate cell to cell communication. Under pathological conditions like glioblastoma (GB), cytonemes transform into ultra-long tumour microtubes (TMs). These filopodia infiltrate through the brain, enwrap neurons and deplete wingless (Wg)/WNT, as a consequence GB progress and neurons undergo synapse loss and degeneration. Thus TMs emerge as a central cellular feature of GB which correlates with a poor prognosis in patients and animal models. Here we describe in a *Drosophila* model for GB the molecular mechanisms behind TMs production, infiltration and maintenance. Glial cells are initially transformed into malignant GB upon EGFR and PI3K pathways constitutive activation, afterwards GB cells establish a positive feedback loop including Wg signalling, JNK and matrix metalloproteases (MMP). In order, Frizzled1 mediates Wg signalling upregulation which activates JNK in GB. As a consequence, MMPs are upregulated and facilitate TMs infiltration in the brain, hence GB TMs network expands and mediate further wingless depletion to close the loop.

## Introduction

Glioblastoma multiforme (GB) is the most frequent and aggressive primary malignant brain tumour with a 3 per 100.000 incidence per year [1]. GB patients’ median survival is 12-15 months, with less than 5% of survival after 5 years [1–4]. The causes of GB are under debate [2], 5% of the patients develop GB after a low grade astrocytoma [5] and the most frequent mutations include gain of function of EGFR (97%) and PI3K/AKT pathways (88%) [6]. The diagnosis, and therefore the treatment of GB, requires a mutations analysis taking into account the high frequency of clones within the same primary GB [7]. Temozolomide (TMZ) has emerged as an effective treatment for GB however, recent discoveries restrict the use of TMZ in GB patients depending on the methylation status of *methylguanine DNA methytransferase (MGMT)* [7]. Moreover, among other mutations, *Isocitrate dehydrogenase (IDH)* define the nature and features of GB [8] together with molecular alterations including 1p/10q deletions and *tumour suppressor protein 53 (TP53)* and *alpha thalassemia/mental retardation (ATRX)* mutations [8, 9]. This genetic and molecular heterogeneity difficult the diagnosis and treatment of this fatal brain tumours.

The recent discovery of an ultra-long tumour microtubes (TMs) network in GB [10] improves our understanding of GB progression and therapy resistance [11]. TMs are actin-based filopodia which infiltrate through the brain and reach long distances within the brain [10]. TMs are required in GB cells to mediate Wingless(Wg)/WNT signalling imbalance among neurons and GB cells. Wg/WNT signal is favoured in GB cells to promote proliferation, at expenses of the neurons which undergo degeneration and cause lethality [12, 13]. The central role of TMs in GB biology has emerged as a fundamental mechanism for GB rapid and lethal progression thus; it is an attractive field of study towards potential GB treatments. However, the molecular mechanisms underlying the expansion of TMs are not well understood and the signalling pathways mediating TM infiltration are still unknown.

Matrix metalloproteases (MMPs) are a family of endopeptidases capable of degrading the extracellular matrix (ECM). Members of the MMP family include the "classical" MMPs, the membrane-bound MMPs (MT-MMPs), the ADAMs (a disintegrin and metalloproteinase; adamlysins) and the ADAMTS (a disintegrin and metalloproteinase with thrombospondin motif). There are more than 20 members in the MMP and ADAMTS family including the collagenases, gelatinases, stromelysins, some elastases and aggrecanases [14]. The vertebrate MMPs have overlapping substrates, they exhibit genetic redundancy and compensation, and pharmacological inhibitors are non-specific. In contrast, there are only two MMP genes in *Drosophila*, MMP1 and MMP2, categorized by their pericellular localization, as MMP1 is secreted and MMP2 is membrane-anchored, suggesting that protein localization was the critical distinction in this small MMP family. Recent reports propose that products of both genes are found at the cell surface and released into media and that GPI-anchored MMPs promote cell adhesion when they are rendered inactive. Moreover, the two MMPs cleave different substrates, suggesting that this is the important distinction within this smallest MMP family [15]. MMPs are upregulated in a number of tumours, including gliomas. MMPs upregulation in GB is associated with the diffuse infiltrative growth and have been proposed to play a role in glioma cell migration and infiltration [16, 17] reviewed in [18]. In consequence, MMPs upregulation in GB is an indicator of poor prognosis [19] and the study of the mechanisms mediated by MMPs is relevant for the biology of GB, and cancer in general.

The Jun-N-terminal Kinase (JNK) pathway has been associated to glial proliferation. GB cells normally activate JNK pathway to maintain the stem-like GB cells, which has become a pharmacological target for the treatment of GB [20]. Moreover, the JNK pathway is the main regulator of MMPs expression and cell motility in different organisms and tissues [21–25].

In this study, we have used the *Drosophila melanogaster* model of glioma [26] to demonstrate that GB cells activate the JNK pathway and upregulate MMPs (MMP1 and 2) expression. MMPs contribute to TMs expansion through the brain and facilitate Frizzled1-mediated Wg/WNT signalling in GB cells. To complete the positive feedback loop, Wg/WNT signalling mediates JNK activation in GB cells to continue with the process. We postulate that the founder mutations in GB (PI3K and EGFR) initiate the process with the expansion of the TMs; afterwards, the system self-perpetuates (TMs-Fz1/Wg-JNK-MMPs-TMs) to facilitate GB progression and infiltration in the brain.

## RESULTS

### MMPs are upregulated in GB

GB cells tumour microtubes (TMs) infiltrate the brain and contribute to tumour progression and associated neurodegeneration [10, 13, 27]. TMs are actin-based structures similar to cytonemes [10, 13, 28] which mediate the exchange of signalling molecules among cells that contribute to GB advancement [13]. The mechanisms mediating TMs formation and progression are not well understood, we propose that TMs infiltration through the brain extracellular matrix (ECM) is mediated by Matrix Metalloproteases (MMPs) activity. To determine if MMPs are expressed in GB, we used specific antibodies and reporters to detect *Drosophila MMP1* or *MMP2* expression. Confocal images show larvae brains stained with anti-MMP1 (Figure 1A-B’’, E, F) or anti-MMP2 (Figure S1A-B’’, E and a GFP reporter shown in F) signal increase in GB compared to control glial cells or to neighbouring neurons. To visualize the cytonemes (WT brain) or the TMs (GB samples) we use the specific marker ihog-RFP. MMPs proteins colocalize with TMs and are preferentially accumulated in the limiting region of GB and healthy brain tissue (Figure 1B-B’’, S1B-B’’ and S1F-F’’). To determine the hierarchy of events between TMs formation and MMPs expression, we attenuated Gap43 expression in GB cells to prevent TMs formation and stained for MMPs. The results show that TMs do not expand upon Gap43 RNAi expression and MMP1 and MMP2 protein levels are comparable to control levels (Figure 1C-C’’, E, F and S1C-C’’, E). Moreover, we reported recently that GB expansion requires Wg signaling, GB cells vampirize Wg from neighbouring neurons via Frizzled1 receptor (Fz1) accumulation in TMs. To determine Wg pathway contribution to MMP1/2 expression, we silenced *Fz1* in GB to reduce Wg-pathway signaling and stained for MMPs. As previously reported, GB cells do not over proliferate; in addition MMP1 and MMP2 are not overexpressed upon *Fz1 RNAi* (Figure 1D-D’’, E, F and S1D-D’’, E). All together the results suggest that *MMP1* and *MMP2* expression depend on TM network formation and Fz1-mediated Wg signalling pathway. These results indicate that GB cells form a TM network which facilitates Wg signalling and then, upregulate MMPs to mediate infiltration of glioma cells in the brain.

**Figure 1:**
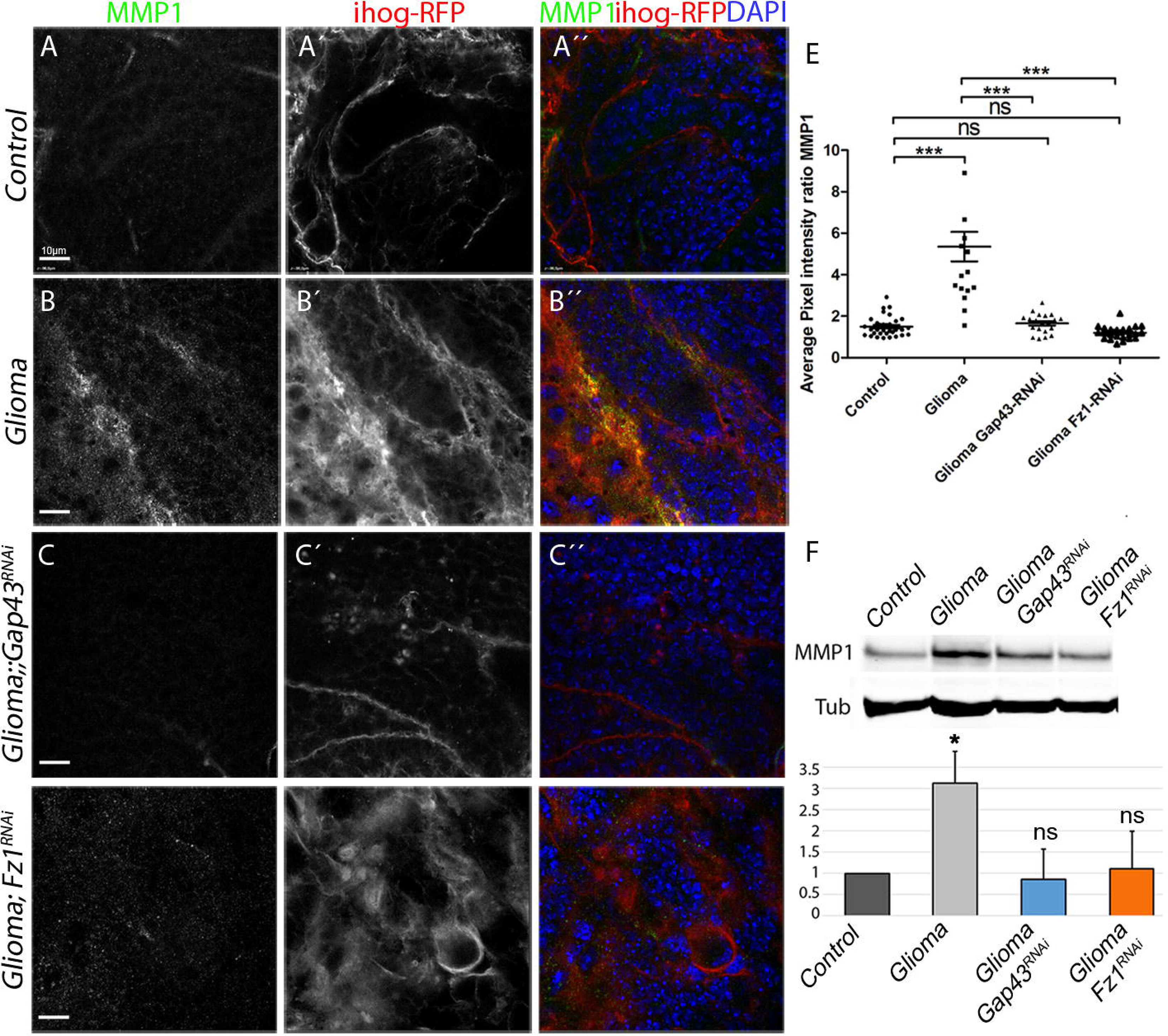
MMP1 is upregulated in GB. Brains from 3rd instar larvae displayed at the same scale. Glia are labeled with *UAS-Ihog-RFP* driven by *repo-Gal4* to visualize active cytonemes/ TM structures in glial cells, and stained with MMP1 (green). (A) MMP1 is homogeneously distributed in control sections, with a slight accumulation in the Ihog+ projections (B) MMP1 accumulates in the TMs and specifically in the projections that are in contact with the neuronal clusters. (C) Inhibition of *Gap43* by *RNAi* in glioma brains restores a normal glial network and MMP1 does not accumulate showing a homogeneous staining along the brain section. (D) Inhibition of *Fz1* by *RNAi* in glioma brains restores a normal MMP1 distribution. MMP1 does not accumulate showing a homogeneous staining along the brain section. Nuclei are marked with DAPI. (E) Quantification of MMP1 staining ratio between ihog^+^ and ihog^−^ domains. (F) Western blot of samples extracted from control, glioma, glioma Gap43-RNAi and Glioma Fz1-RNAi larvae showing changes in the amount of MMP1 or protein. Error bars show S.D. * P<0.01, ** P<0.001, *** P<0.0001 or ns for non-significant. Scale bar size is indicated in this and all figures.

### PI3K or EGFR individually do not stimulate MMPs expression

To determine the epistatic relations behind MMPs expression, we studied single gene modifications related to GB and the effect on MMPs. We expressed the constitutively active forms of PI3K (dp110) and EGFR (TOR-DER^CA^) in glial cells marked with ihog-RFP. In both cases we did not observe an upregulation of MMP1 or the formation of TMs (Figure S2A-B’’ and D). PI3K and EGFR pathways converge in *dMyc* expression [5], thus we tested *dMyc* upregulation in glial cells as a candidate to upregulate MMPs and form a TMs network. However, we did not observe any significant change in MMPs expression or TM expansion upon *dMyc* upregulation (Figure S2C-C’’ and D). Taking these results together, even though Myc is a convergent point of EGFR and PI3K pathways, it is not sufficient to reproduce the features of glioma. These results suggest that both PI3K and EGFR together are necessary to activate a downstream pathway responsible for the expansion of glial projections and MMP1 accumulation in glial transformed cells, but Myc expression is not the cause of these phenotypes.

The results show that Fz1 is necessary to upregulate MMPs (Figure 1D-D’’ and S1D-D’’) upon GB induction. To conclude if Fz1 is sufficient to trigger MMP upregulation we overexpressed Fz1 in WT glial cells (Figure 2A-E and S1G-I). The results show MMP1 and MMP2 upregulation in glial cells upon Fz1 overexpression as compared to control samples (Figure 2A-E and S1G-I). These results suggest that Wg pathway is necessary and sufficient to activate MMP expression in glial cells.

**Figure 2:**
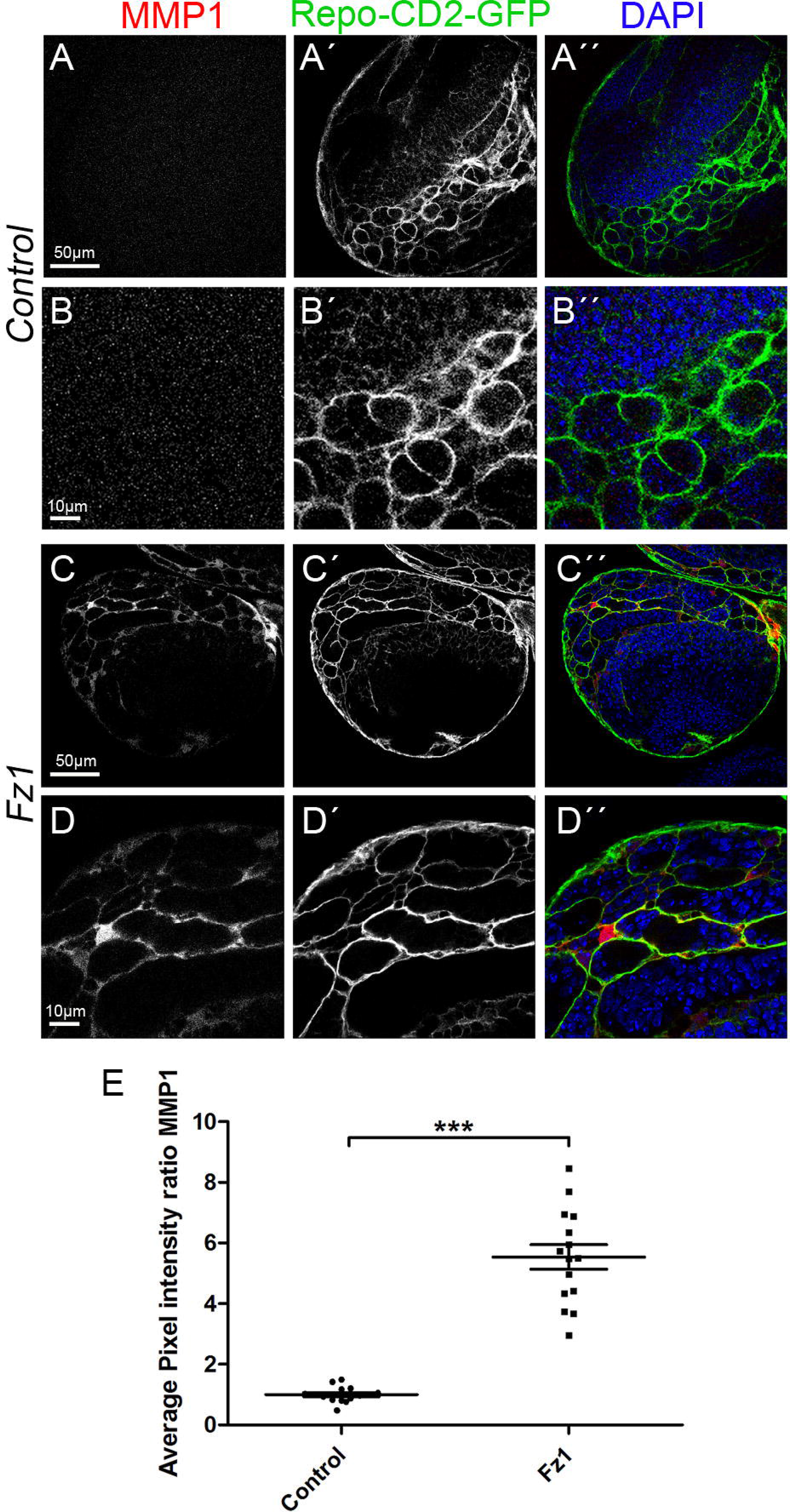
Wg pathway is necessary and sufficient to activate MMP expression in glial cells. Brains from 3rd instar larvae displayed at the same scale. Glial cell bodies and membranes labelled in green (CD8-GFP) driven by *repo-Gal4* to the glial cells and stained with MMP1 (red). (A-B) MMP1 is homogeneously distributed in control sections. (C-D) MMP1 accumulates in the glial cells upon Fz1 overexpression. (E) Quantification of MMP1 staining ratio between GFP^+^ and GFP^−^ domains. Nuclei are marked with DAPI. Error bars show S.D. * P<0.01, ** P<0.001, *** P<0.0001 or ns for non-significant. Scale bar size is indicated in this and all figures.

### JNK activation in glioma

JNK is upregulated in a number of tumours including GB and it has been related to glioma malignancy [29–32]. Moreover, JNK is a target for specific drugs in combination with temozolomide treatments as it was proven to play a central role in GB progression [20, 33–35]. However, little is known about the molecular mechanisms underlying JNK activation in glioma cells and the functional consequences for GB progression.

To confirm JNK pathway activation in GB cells, we used two independent reporters. *puc-LacZ* monitors the transcriptional activation of the downstream JNK target *puckered [36, 37]* in addition, *Tre-GFP* confer transcriptional activation in response to JNK signalling [38–40]. GB cells show a progressive upregulation of *puc-LacZ* (Figure 3A-D) and Tre-GFP reporters (Figure 3E-F, J) compared to control samples, indicating that JNK pathway is activated in GB cells. Moreover, to determine if JNK activation depends on TMs formation or Wg pathway activity, we silenced *Gap43* or *Fz1* respectively and monitored *Tre-GFP* and quantified the number of Repo^+^ cells in GB brains (Figure 3G-J). The results show that JNK pathway depends on the formation of the TMs network as well as the presence of Fz1 receptor in GB cells.

**Figure 3:**
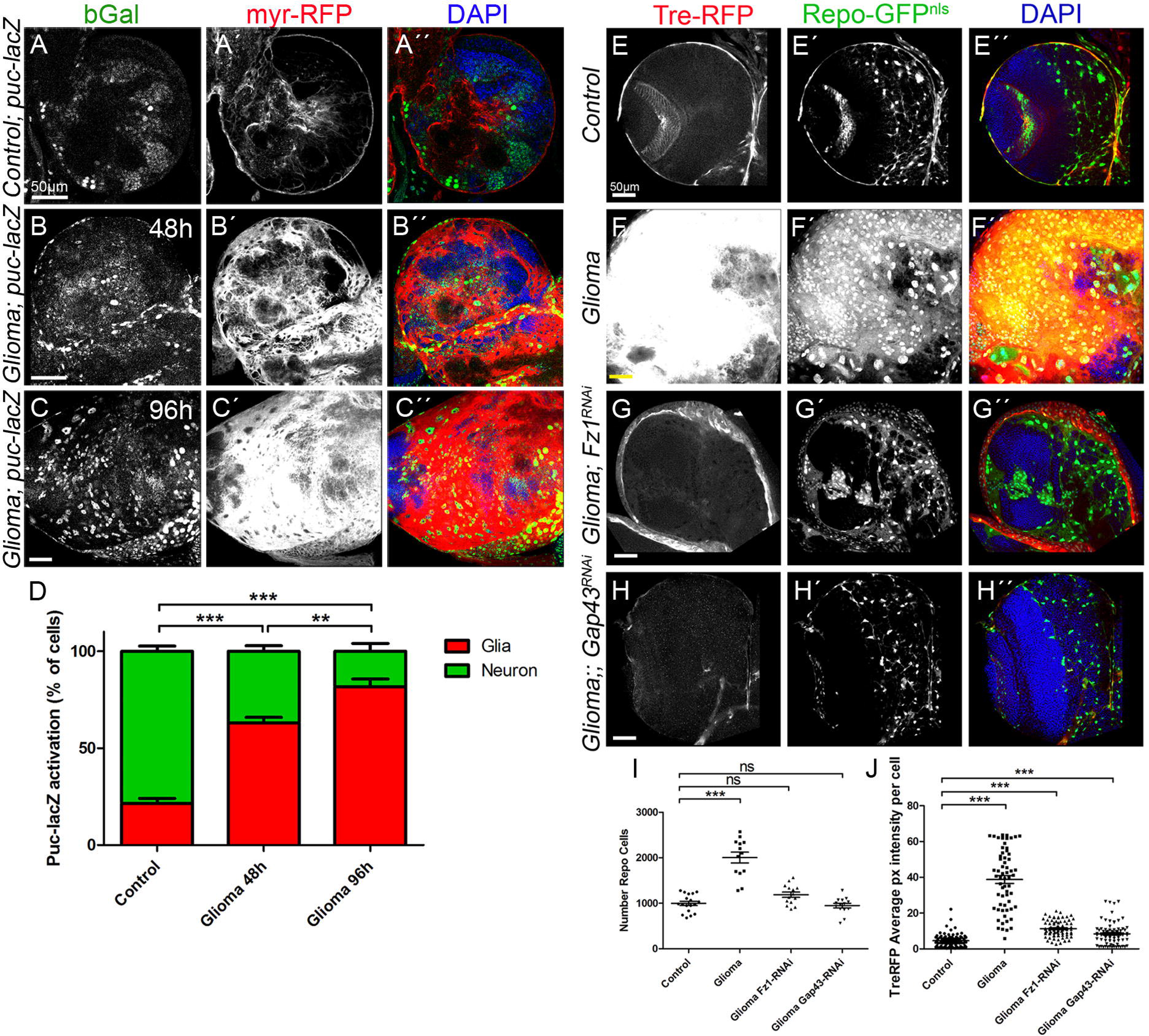
JNK signaling pathway activation in glioma. Larval brain sections from 3rd instar larvae displayed at the same scale. Glial cell bodies and membranes are labeled with *UAS-myr-RFP* (red) driven by *repo-Gal4* (A-C) JNK signaling pathway reporter puc-lacZ in control, glioma 48h and glioma 96h After Tumour induction, shows activation of the pathway mostly in neurons in control samples (A) and then shows a progressive activation in glial transformed cells (B-C). (D) Quantification of the % of cells with Puc-lacZ activation in glial cells and neurons domains. Glial cell nuclei are labeled with *UAS-GFP*^*NLS*^ (Green) driven by *repo-Gal4.* (E-H) JNK signaling pathway reporter TRE-RFP in control, glioma, glioma Gap43-RNAi and Glioma Fz1-RNAi brain sections. The number of glial cells is quantified in (I) and the TRE-RFP average pixel intensity per glial cells is quantified in (J). Nuclei are marked with DAPI. Error bars show S.D. * P<0.01, ** P<0.001, *** P<0.0001 or ns for non-significant. Scale bar size is indicated in this and all figures.

### Grindelwald receptor mediates GB progression

Next we decipher the molecular basis of JNK pathway activation in GB cells. eiger (egr) is the ligand for JNK-grnd pathway, to determine the localization of we used a egr-GFP protein fusion and monitored GFP signal in confocal images. The results show that GFP signal is localized in the neurons that contact with glial cells and the egr-GFP signal is higher in GB samples (Figure S3A-E). Next we monitored the expression pattern of the JNK receptor Grindelwald (grnd), we used a specific antibody in GB and control samples (Figure S3F-H). The results show that grnd protein accumulation in glial membranes is increased in GB brains. Taking these data together, GB cells increase the amount of grnd and egr protein is upregulated in the tissues surrounding GB cells.

To study the contribution of grnd to GB progression, we attenuated *grnd* expression with a *UAS-grnd RNAi* expressed in glial cells under the control of *repo-Gal4*. The results show that *grnd* knockdown does not reduce the number of glial cells during development (Figure 4A, C, G) but prevents the increase of glial cells upon GB induction (Figure 4B, D, G). To further validate the results, we analysed the number of glial cells in a mutant background for *grnd* (see materials and methods). The results reproduced the previous observations, the number of glial cells was not affected during development but in the case of GB brains, glial cell number was not increased (Figure 4E-G). Thus, grnd receptor is dispensable for glial development and necessary in GB cells for tumour advance.

**Figure 4:**
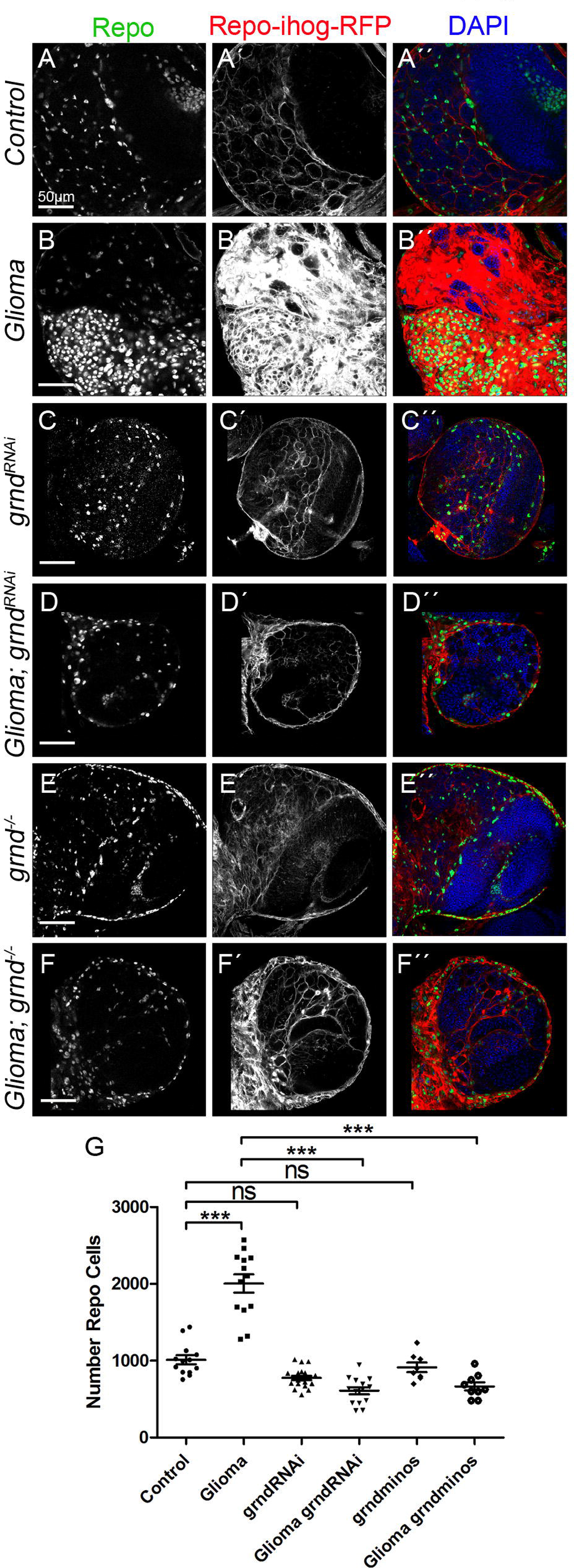
TNF receptor Grindelwald mediates GB progression. Brains from 3rd instar larvae displayed at the same scale. Glia are labeled with *UAS-Ihog-RFP* driven by *repo-Gal4* to visualize active cytonemes/ TM structures in glial cells, and stained with Repo (green) in the following genotypes (A-F) control, glioma, grnd-RNAi, glioma grnd-RNAi, grnd^−/−^ and Glioma grnd^−/−^ brain sections. The number of Repo^+^ cells is quantified in (G). Nuclei are marked with DAPI. Error bars show S.D. * P<0.01, ** P<0.001, *** P<0.0001 or ns for non-significant. Scale bar size is indicated in this and all figures.

To further validate the contribution of JNK pathway in GB progression, we modified JNK pathway expressing a dominant negative form of the effector Bsk (Bsk^DN^) in glial cells, or inducing GB in a *egr* mutant background (egr−/−), or expressing the external domain of grnd which causes a dominant negative effect (grnd extra) in GB cells. In all three situations, JNK pathway disruption in GB cells prevent the increase of glial cells number as compared to controls (Figure S4). Moreover, these modifications also cause a reduction in the number of wt glial cells compared to controls suggesting that JNK signalling pathway is required for normal glial development and GB progression.

### grnd mediates Wg pathway activation in GB

Wg pathway activity in glial cells is necessary for GB development, as previously described [1]. GB cells accumulate Fz1 receptors which mediate Wg signalling in the tumour at expenses of neuronal depletion of Wg and, as a consequence, results in neurodegeneration. This accumulation of Fz1 in TMs depends on Gap43. To establish the relation between JNK activation in GB and Wg pathway, we analysed the contribution of *grnd* in Fz1 distribution (Figure 5), and the activity of Wg pathway (armadillo) in GB brains upon *grnd* depletion (Figure 6).

**Figure 5:**
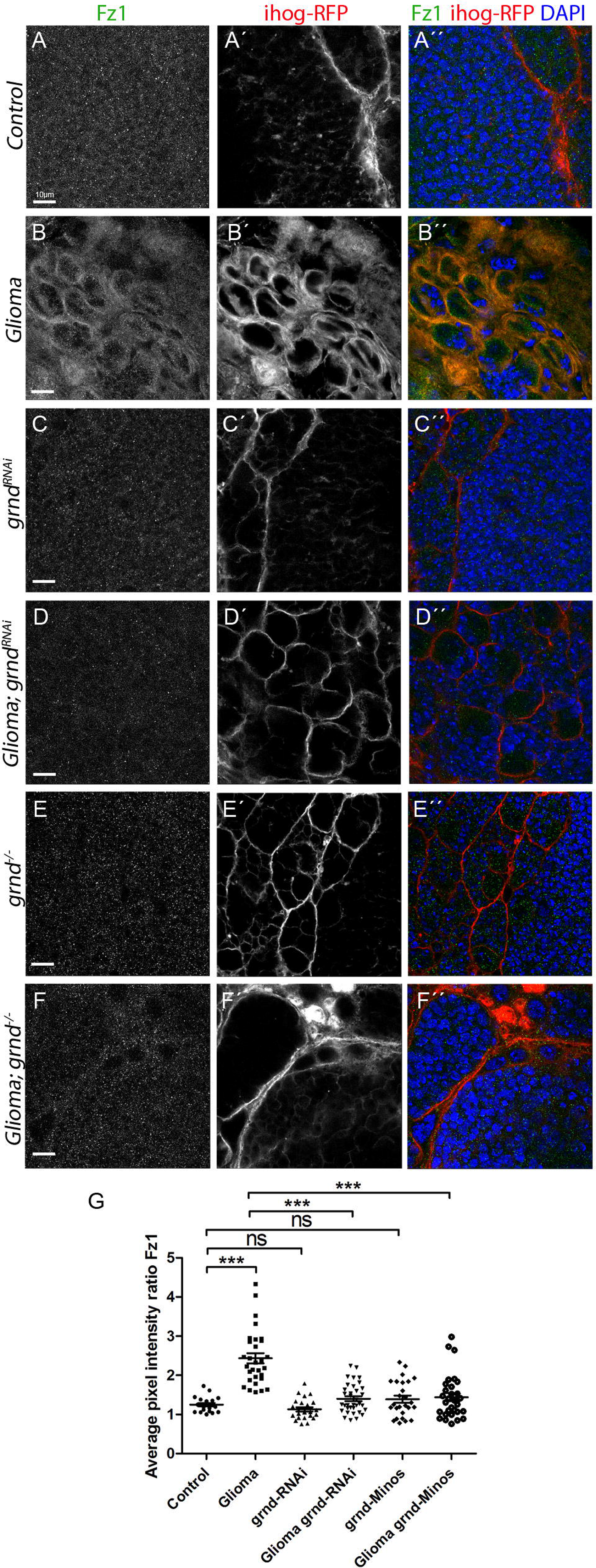
JNK pathway is necessary to localize Fz1 in the TMs and promote GB progression. Brains from 3rd instar larvae displayed at the same scale. Glia are labeled with *UAS-Ihog-RFP* driven by *repo-Gal4* to visualize active cytonemes/ TM structures in glial cells, and stained with Fz1 (green) in the following genotypes (A-F) control, glioma, grnd-RNAi, glioma grnd-RNAi, grnd^−/−^ and Glioma grnd^−/−^ brain sections. Fz1 average pixel intensity staining quantification ratio between ihog^+^ and ihog^−^ domains (G). Nuclei are marked with DAPI. Error bars show S.D. * P<0.01, ** P<0.001, *** P<0.0001 or ns for non-significant. Scale bar size are indicated in this and all figures.

**Figure 6:**
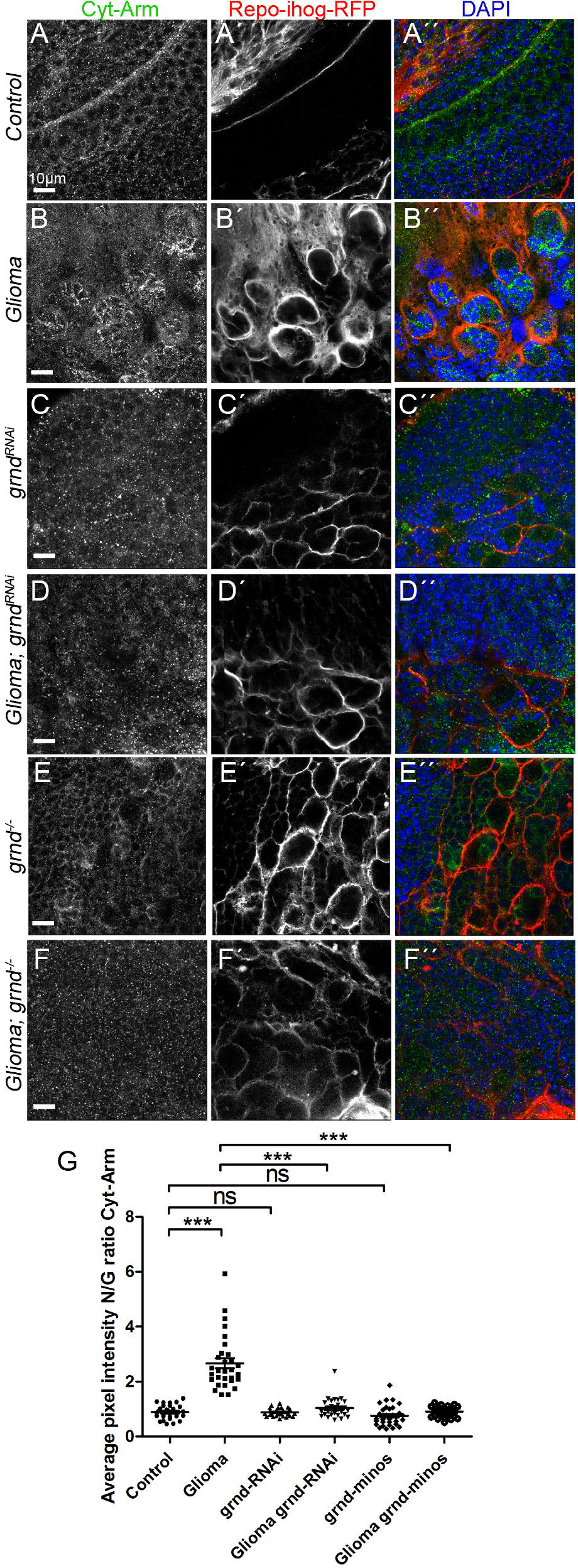
grnd mediates Wg pathway activation in GB. Brains from 3rd instar larvae displayed at the same scale. Glia are labeled with *UAS-Ihog-RFP* driven by *repo-Gal4* to visualize active cytonemes/ TM structures in glial cells, and stained with Cyt-Arm (green) in the following genotypes (A-F) control, glioma, grnd-RNAi, glioma grnd-RNAi, grnd^−/−^ and Glioma grnd^−/−^ brain sections. (A) Cytoplasmic-Armadillo (Cyt-Arm) is homogeneously distributed in control sections. (B) In glioma brains Cyt-Arm accumulates in the neurons cytoplasm where it is inactive. (C-F) Knockdown or knockout of *grnd* in both normal or glioma brains shows a normal glial network and Cyt-Arm does not accumulate showing a homogeneous distribution similar to the control. Cyt-Arm average pixel intensity quantification for Neuron/Glia ratio between ihog^−^ and ihog^+^ domains (G). Nuclei are marked with DAPI. Error bars show S.D. * P<0.01, ** P<0.001, *** P<0.0001 or ns for non-significant. Scale bar size is indicated in this and all figures.

Fz1 is homogeneously distributed through the brain under normal conditions (Figure 5A-A’’, G) but, under GB conditions, Fz1 is accumulated in the TMs surrounding the neurons (Figure 5B-B’’, G). The knockdown of *grnd* expression in glial cells (Figure 5C-C’’, G) does not affect Fz1 distribution and in GB cells (Figure 5D-D’’, G), *grnd* RNAi restores Fz1 distribution to a normal situation. To further confirm the effect of *grnd* knockdown on glia and GB cells, we performed identical experiments in a *grnd* mutant background. Again Fz1 distribution requires *grnd* expression in GB cells to accumulate around neurons (Figure 5E-G). In addition, we blocked JNK pathway downstream grnd with a dominant negative form of Bsk (Bsk^DN^) [41] in GB cells (Figure S5). The results show that JNK pathway is necessary to localize Fz1 in the TMs and promote GB progression

Next we studied the activation of Wg signalling pathway in glia and GB cells. Armadillo (Arm) is the orthologue of mammalian β-catenin which translocate to the nuclei to activate transcription of target genes upon Wg-Fz1 signalling [13, 42]. We used an specific antibody (Cyt-Arm) which detects the inactive form of Arm [13, 42] to monitor the activity of the pathway. Control brains show homogeneous signal for Cyt-Arm (Figure 6A-A’’, G) but, as a consequence of Fz1 distribution, GB cells activate Wg pathway as represented by the reduction of Cyt-Arm signal (Figure 6B’B’’, G). *grnd RNAi* or *grnd* mutant background prevent the imbalance of Wg signalling among GB cells and neurons and restores Wg signalling to a control situation (Figure 6C-G), similar to previous results obtained for Fz1 distribution,

In addition, we analysed the contribution of JNK pathway to the progression of TMs in GB. We used the reporter ihog-RFP to visualize cytonemes in wt glial cells and TMs in GB. The confocal images show that upon downregulation of *grnd* expression or the inhibition of JNK pathway by *Bsk^DN^* in GB cells, the formation of TMs necessary for GB development does not occur (Figure 5 and Figure 6). These results suggest that JNK pathway activation mediated by the receptor grnd is a requirement for TMs formation. As a consequence of the prevention of TMs network development, GB cells do not localize Fz1 in the areas of contact with neurons and cannot mediate Wg depletion. This leads to a reduction in the proliferation of GB cells. Thus the results indicate that grnd activity mediates JNK signalling to form the TMs network in GB necessary to facilitate tumour progression.

### JNK triggers MMPs expression in GB

The mechanisms for TMs progression and GB cells infiltration through the brain are not well understood. GB cells project TMs which cross the extracellular matrix (ECM) and break in the brain to reach territories distant from the original GB site. We propose that JNK activity in GB mediate the production of matrix metalloproteinases (MMPs) to facilitate ECM digestion and, as a consequence, permit TMs infiltration through the brain.

To validate this hypothesis, we first determined if MMPs expression in GB (Figure 1, Figure 7A-B’’) depends on grnd and JNK pathway signalling. We generated a GB and silenced JNK pathway with Bsk^DN^ (Figure 7C-C’’) or silenced *grnd* expression (Figure 7D-E’’). The quantification of MMP1 signal showed that GB upregulation of MMP1 is prevented after JNK pathway silencing (Figure 7F). As occurs in epithelial cells, MMP1 is a target of JNK pathway also in GB cells.

**Figure 7:**
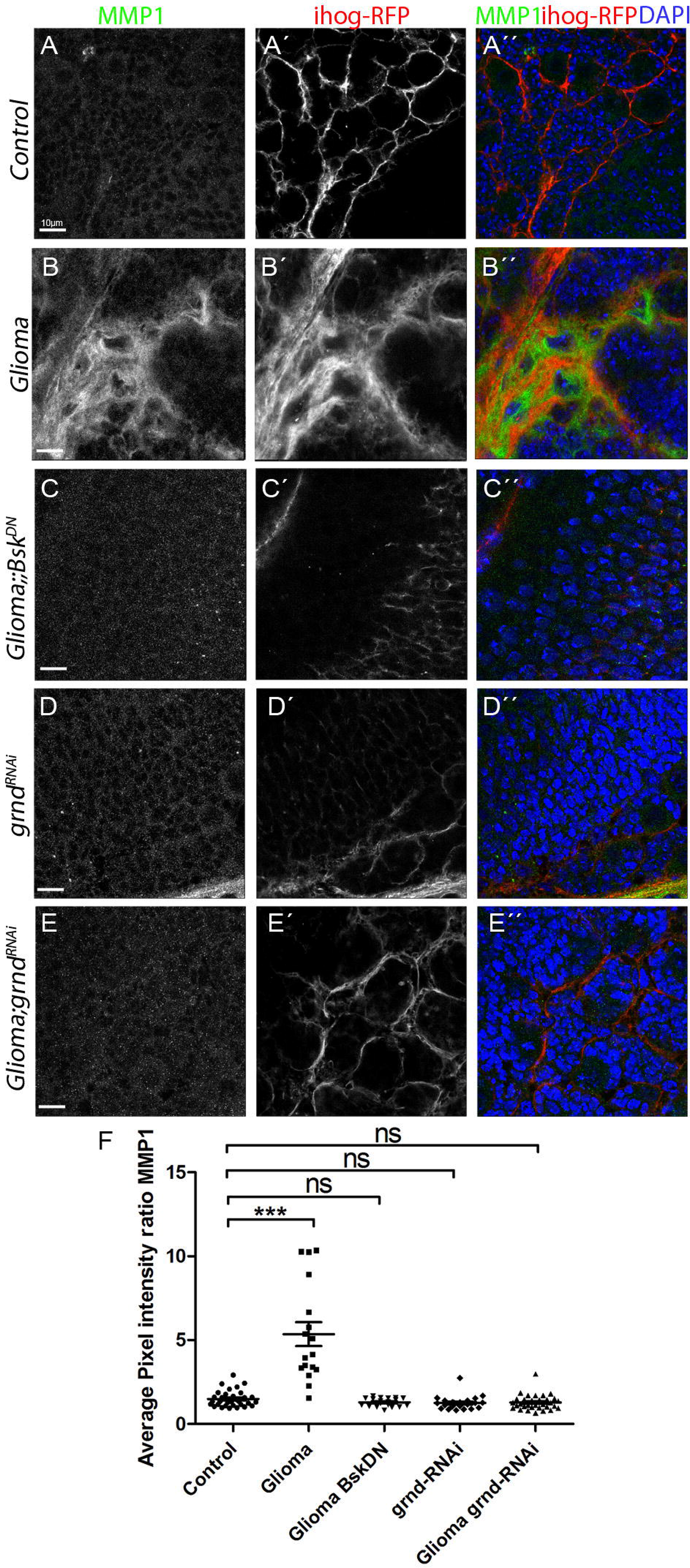
JNK triggers MMPs expression in GB. Brains from 3rd instar larvae displayed at the same scale. Glia is labelled with *UAS-Ihog-RFP* driven by *repo-Gal4* to visualize active cytonemes/ TM structures in glial cells, and stained with MMP1 (green). (A) MMP1 is homogeneously distributed in control sections (B) MMP1 accumulates in the TMs and specifically in the projections that are in contact with the neuronal clusters. (C-E) Blocking JNK pathway by using a *UAS-bsk*^*DN*^ or *UAS-grnd-RNAi* in glioma brains restores a normal glial network and MMP1 does not accumulate showing a homogeneous staining along the brain section. (F) Quantification of MMP1 staining ratio between ihog^+^ and ihog^−^ domains. Error bars show S.D. * P<0.01, ** P<0.001, *** P<0.0001 or ns for non-significant. Scale bar size are indicated in this and all figures.

### TMs expansion requires MMPs

Matrix metalloproteinases (MMPs) are extracellular enzymes responsible for the degradation of the extracellular matrix; cancer cells produce MMPs to facilitate tumour progression and invasiveness. MMPs are upregulated in human GBM cell-lines and biopsies as compared with low-grade astrocytoma (LGA) and normal brain samples [43, 44]. In particular, among the 23 MMPs present in humans, MMP9, MMP2 and MMP14 are directly implicated in growth and invasion of GB cells [45].

WNT induces MMPs expression during development and cancer [46–50] associated to cell migration and metastasis. Specifically in human GBs, MMP2 expression and their infiltrative properties correlate with Wnt5 [51, 52] and MMP9 is upregulated upon EGFR activity [53]. To elucidate if MMP expression is related to the development of the network as a mechanism to cooperate in tumour infiltration, we analyzed MMPs expression in *Drosophila* gliomas. There are two orthologues to human MMPs in *Drosophila*, MMP1 and MMP2.

We have determined that MMP1 is upregulated in glioma tissue induced by Fz1 receptor (Wg pathway) accumulation in TMs (Figure 1) and JNK activation (Figure 7).

To determine the contribution of MMPs to Fz1 receptor accumulation and Wg vampirization [13] we carried out functional studies. But first, *UAS-MMP1* and *UAS-MMP2 RNAi* tools [54] were validated in epithelial tissues (Figure S6). *MMP1* or *MMP2 RNAi* were expressed in GFP-marked cells (see material and methods) and stained with specific MMP1 or MMP2 antibodies. Both RNAi tools reduced significantly the signal for MMP1 or MMP2 staining.

Afterwards we silenced *MMP1* or *MMP2* in GB and stained the brain samples for Fz1 and Wg (Figure 8A-G S7A-G). Confocal images from larval brains show that Fz1 distribution in control brains (Figure 8A-A’’) and GB (Figure 8B-B’’). The results show that *MMP1* or *MMP2 RNAi* does not disrupt Fz1 brain distribution (Figure 8C-C’’, E-E’’, G) nor glial cell cytoneme network (Figure 8H). In line with previous reports [13] Fz1 is accumulated in GB cells membranes and this is halted by *MMP1* or *MMP2* knockdown (Figure 8D-D’’, F-F’’, G). Moreover *MMP1* or *MMP2* interference prevents TMs network formation and expansion in GB (Figure 8D’, F’ and H).

**Figure 8:**
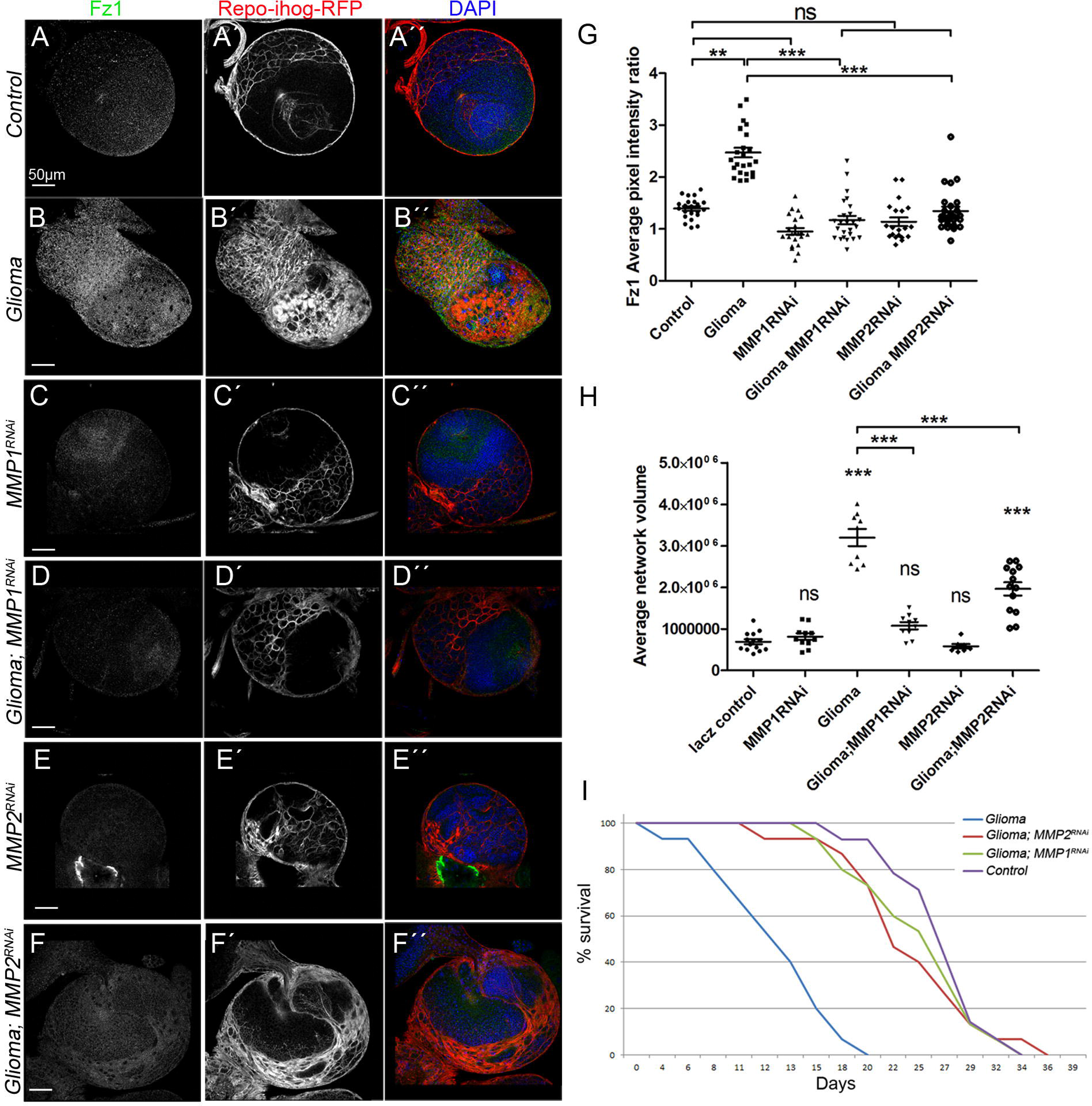
TMs expansion requires MMPs. Brains from 3rd instar larvae displayed at the same scale. Glia are labeled with *UAS-Ihog-RFP* driven by *repo-Gal4* to visualize active cytonemes/ TM structures in glial cells, and stained with Fz1 (green) in the following genotypes (A-F) control, glioma, MMP1-RNAi, glioma MMP1-RNAi, MMP2-RNAi and glioma MMP2-RNAi brain sections. (G) Quantification of Fz1 average pixel intensity staining ratio between ihog^+^ and ihog^−^ domains. (H) Quantification of glial/glioma network volume expansion. Nuclei are marked with DAPI. (I) Survival curve of adult control, glioma, glioma MMP1-RNAi and glioma MMP2-RNAi flies after a number of days of glioma induction and progression. Error bars show S.D. * P<0.01, ** P<0.001, *** P<0.0001 or ns for non-significant. Scale bar size is indicated in this and all figures.

In addition, Wg accumulation in GB membranes (Figure S7 B, D, F, and G) is prevented by *MMP1* or *MMP2* attenuation. Moreover, *MMP1* or *MMP2 RNAi* does not affect Wg distribution in control samples (Figure S7A, C, E, and G). Next, to determine the contribution of MMP1 and MMP2 expression in GB to Wg signalling we downregulated MMP1 or MMP2 in GB samples using a *Repo-Gal4* driver and a *UAS-ihog-RFP* to mark the glial network and stained for Cyt-Arm. Control brains show homogeneous signal for Cyt-Arm (Figure S7H-H’’, N) but, as a consequence of Fz1 distribution, GB cells activate Wg pathway as represented by the reduction of Cyt-Arm signal (Figure S7I-I’’, N). Similar to previous results obtained for Fz1 distribution, *MMP1 or MMP2 RNAi* prevent the imbalance of Wg signalling among GB cells and neurons and restores Wg signalling to a control situation (Figure S7J-N), similar to previous results obtained for Fz1 distribution.

To determine the contribution of MMPs to the lethality caused by GB, we downregulated *MMP1* or *MMP2* in GB samples and analyzed survival. To avoid developmental defects caused by *MMP1* or *MMP2* attenuation, we did the survival experiments after metamorphosis, in adults. We used the Gal80^TS^ system to silence the binary expression system UAS/Gal4 during development, and switched it on 4 days after eclosion. Glioma lethality was rescued by *MMP1* or *MMP2* down regulation (Figure 8I) suggesting that MMPs are required for GB lethality. All the data together suggest that MMPs facilitate infiltration and tumour invasiveness in the brain, mediate Wg pathway imbalance, GB cells increase and TMs projection which induce premature death.

GB causes neurodegeneration and neurological symptoms in patients [55]. We have previously described that in GB brains TMs expansion and therefore the accumulation of Fz1 in the projections is having an effect on the neighbouring neurons that accumulate inactive Arm [13]. Moreover, it has been reported that reduction in Wg signalling causes reduction of synapse number, which is one of the initial events in neurodegeneration [56–59]. We wondered if MMPs role in GB would reduce the synapse number in our animal model. We specifically inhibited *MMP1* or *MMP2* in GB and quantified the number of active zones in neuromuscular junctions. The results showed a significant reduction in the number of synapses in the neurons upon GB induction compared with the control (Figure 9A-B). However, upon *MMP1* or *MMP2* knockdown in GB, neurons are protected and the number of active zones is rescued (Figure 9D, F, G).

**Figure 9:**
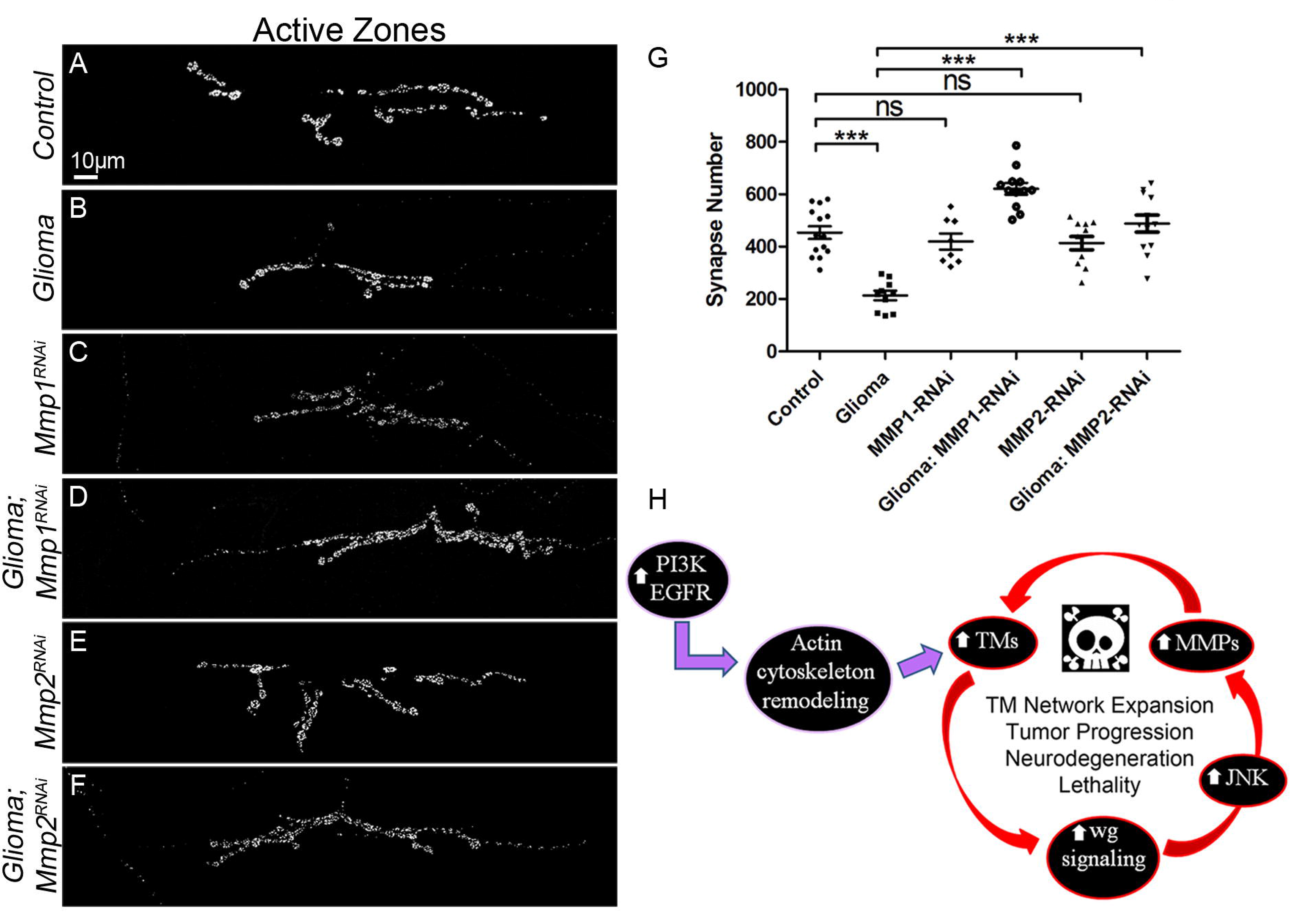
MMPs upregulation in glioma are responsible for the synapse loss in the neurons. Neurons from the larval neuromuscular junction are stained with Nc82 (brp) showing the synaptic active Zones in grey. (A-F) Upon glioma induction (B) the number of synapses (grey) is reduced when compared with the control (A). The number of synapses is restored upon knockdown of *MMP1* or MMP2 (D, F).The quantification of the number of synaptic active zones in all genotypes is shown in (G). (H) Model: Glial cells are initially transformed into malignant GB upon EGFR and PI3K pathways constitutive activation, afterwards GB cells establish a positive feedback loop including TMs, Wg signalling, JNK and matrix metalloproteinases (MMPs). Initial stimulation of actin cytoskeleton remodelling via EGFR/PI3K enables initial expansion of TMs, as a consequence Frizzled1 receptor accumulation in TMs mediates neuronal Wg depletion and Wg signalling upregulation in the GB cells which activates JNK in GB. As a consequence, MMPs are upregulated and facilitate further TMs infiltration in the brain, hence GB TMs network expands and mediate further wingless depletion to close the loop.

Taking all these data together, we showed that MMPs mediate GB progression and neurodegeneration. Further, *MMPs* silencing in GB cells is sufficient to rescue life spam in adult flies. These results bring a novel positive feedback loop among Wg pathway, JNK and MMPs which contribute to GB development (Figure 9H), thus opening novel avenues for GB adaptation mechanisms and potential pharmacological targets.

## Discussion

Activating mutations for EGFR and PI3K pathways are the most frequent initial signals in GB. However, the attempts to treat GB reducing the activation of these pathways have so far been limited by acquired drug resistance. The current tendencies suggest that a multiple approach is required to obtain a more positive result [60–64]. GB cells show a high mutation rate and usually present more than two sub-clones within the same patient and from the same primary tumour [65, 66].

Here we describe that JNK signalling regulates MMPs expression in GB, which is required for TMs network formation and infiltration. As a consequence, Wg pathway responds to JNK and TMs expansion, these three events conform a regulatory positive feedback loop in GB progression (Figure 9H). Grnd is the JNK receptor upregulated in GB cells, it binds to the ligand egr and activates JNK pathway. GFP fusion protein showed that egr is produced in the surrounding neurons and accumulated in the border of GB and healthy neuronal tissue. Therefore, here is another example of neuron-glia molecular interaction which mediates the physiological status of the brain and the evolution of GB. It is of interest to unravel the regulatory mechanisms that mediate *egr* expression and secretion in neurons as a potential modulator for brain tumour advance.

Frizzled receptors mediate matrix metalloproteinase (MMP) 2 and MMP9 expression in different scenarios. Blocking WNT signaling, or MMP activity, reduces T cell migration through the basement membrane in vitro and into inflamed skin in vivo. MMP promoters respond to WNT signalling through tandem TCF sites and mediate T cell extravasation [67]. MMP9 is a key molecular effector, downstream of HIF-1α and WNT activation, responsible for increased rates of neural stem cells proliferation and migration in hypoxia [68]. In addition, *Mmp7* (also known as matrilysin) is a target gene of the Wnt signalling pathway in lung epithelial cells and is known to be a key mediator of pulmonary fibrosis [69]. Moreover, MMPs expression is associated to GB invasion, growth and angiogenesis [70, 71]. MMP2 and MMP9 co-silencing in combination with temozolomide treatment emerge as promising in the treatment against GB [72]. Furthermore, phase II clinical trials using the broad-spectrum MMP-inhibitor, marimastat, in conjunction with temozolomide has shown encouraging results [73]. Thus, the evidences suggest a central role for WNT pathway and specific MMPs in GB and emerge as promising targets for potential treatments [43].

As a result, the tandem JNK-Wg-TMs trigger the expression of *MMPs* in *Drosophila* GB cells. Fly MMP1 and MMP2 degrade the ECM and, in the brain facilitate the infiltration of TMs. MMPs are upregulated in human patients [43] as well as in *Drosophila* GB model. Besides, MMPs expression attenuation reduces the volume occupied by TMs and Wg signalling with the concomitant consequences: reduction of GB progression, prevention of neurodegeneration (synapse loss) and lethality. Consequently, MMPs are part of the positive feedback loop in GB and mediate the equilibrium among Wg signalling, JNK and TMs expansion.

MMPs have been a field of interest for GB for more than a decade, numerous studies correlate MMPs expression with a poor prognosis [43, 74] but the precise mechanisms mediating the cellular impact of MMPs and the regulation of *MMPs* expression has not been elucidated. We propose that GB cells become addicted to signals independent of the founder mutations (PI3K and EGFR), in this case, the positive feedback loop formed by Wg pathway, JNK, MMPs and the TMs (Figure 9H). As a consequence, treatments tackling EGFR of PI3K failed to success as GB cells rely of other inputs. This particular signalling loop is required for GB cells to progress but is dispensable for normal glia development which puts these discoveries as targetable features for GB treatments.

Finally, the specific targets of MMPs will be of interest to determine which ECM components are essential to prevent TMs expansion. There are two MMPs in *Drosophila* (MMP1 and MMP2) with 10 and 3 isoforms respectively, MMP1 and MMP2 have been classically differentiated by their extracellular or membrane associated localization, however this concept is currently under debate [15]. It is proposed that each MMP has particular targets from the ECM which are sensible of degradation, recent classifications have brought light on the specificity of each protease and the different substrates [43]. The results show that the attenuation of either MMP1 or MMP2 prevents GB progression, thus the potential substrates and the functional implications for GB need to be established in future studies.

## Acknowledgements

We thank Professor Alberto Ferrús and anonymous reviewers for critiques of the manuscript and for helpful discussions. We are grateful to R. Read, I. Guerrero, P. Leopold, C. Klambt, JP. Vincent, E. Martín-Blanco, K. Broadie, E. Moreno, the Vienna *Drosophila* Resource Centre, the Bloomington *Drosophila* stock Centre and the Developmental Studies Hydridoma Bank for supplying fly stocks and antibodies, and FlyBase for its wealth of information. We acknowledge the support of the Confocal Microscopy unit and Molecular Biology unit at the Cajal Institute for their help with this project. MP holds a fellowship from the Juan de la Cierva program IJCI-2014-19272 and SCT holds a contract from the Ramón y Cajal program RYC-2012-11410 from the Spanish MICINN. Research has been funded by grant BFU2015-65685P. Authors declare no conflicts of interest.

## Experimental Procedures

### Fly stocks

Flies were raised in standard fly food at 25ºC. Fly stocks from the Bloomington stock Centre: *UAS-GFP*^*nls*^ (BL4776), *UAS-lacZ* (BL8529), *UAS-myr-RFP* (BL7119), *UAS-Gap43-RNAi* (BL29598), *repo-Gal4* (BL7415), *UAS-CD8-GFP* (BL32186), *tub-gal80*^*ts*^ (BL7019), *Repo-lexA (a gift from C Klambt), lexAop-CD2-GFP* (BL32205), *UAS-bsk*^*DN*^ (BL9311), egr-GFP (BL66381), *egr*^*[MI15372]*^ (BL59754), *en-Gal4, UAS-GFP (from BL25752), MMP2-GFP* (BL60512). Fly stocks from the Vienna *Drosophila* Resource Centre: *UAS-fz1-RNAi* (v105493), *UAS-mmp1-RNAi* (v101505), *UAS-mmp2-RNAi* (v107888), *lexAop-Fz1 [13], UAS-dEGFR*^*λ*^, *UAS-PI3K92E* (dp110^CAAX^) (A gift from R. Read), *UAS-ihog-RFP* (a gift from I. Guerrero), TRE-RFP-1b (a gift from J.P. Vincent) *and puc-lacZ* (a gift from E. Martín-Blanco), *UAS-grnd-RNAi*, *grnd*^*MINOS*^ *and UAS-grnd*^*EXTRA*^ (a gift from P. Leopold) [75], *UAS-dmyc* (a gift from E. Moreno) [76], *UAS-TOR-DER*^*CA*^ [77].

### *Drosophila* glioblastoma model

The most frequent genetic lesions in human gliomas include mutation or amplification of the Epidermal Growth Factor Receptor (EGFR) gene. Glioma-associated EGFR mutant forms show constitutive kinase activity that chronically stimulates Ras signaling to drive cell proliferation and migration [78, 79] Other common genetic lesions include loss of the lipid phosphatase PTEN, which antagonizes the phosphatidylinositol-3 kinase (PI3K) signaling pathway, and mutations that activate PI3KCA, which encodes the p110a catalytic subunit of PI3K ^98-99^. Gliomas often show constitutively active Akt, a major PI3K effector. However, EGFR-Ras or PI3K mutations alone are not sufficient to transform glial cells. Instead, multiple mutations that coactivate EGFR-Ras and PI3K/Akt pathways are required to induce a glioma [80]. In *Drosophila*, a combination of EGFR and PI3K mutations effectively causes a glioma-like condition that shows features of human gliomas including glia expansion, brain invasion, neuron dysfunction, synapse loss and neurodegeneration [26, 81, 82]. Moreover, this model has proved to be useful in finding new kinase activities relevant to glioma progression [83] To generate a glioma in *Drosophila* melanogaster adult flies, the Gal4/UAS system [84] was used as described above (*repo*-Gal4>UAS-*EGFRλ*,UAS-*dp110*. To restrict the expression of this genetic combination to the adulthood, we used the thermo sensitive repression system Gal80^TS^. Individuals maintained at 17ºC did not activate the expression of the UAS constructs, but when flies were switched to 29ºC, the protein Gal80^TS^ changed conformation and was not longer able to bind to Gal4 to prevent its interaction with UAS sequences, and the expression system was activated.

### Antibodies for Immunofluorescence

Third-instar larval brains, were dissected in phosphate-buffered saline (PBS), fixed in 4% formaldehyde for 30min, washed in PBS + 0.1 or 0.3% Triton X-100 (PBT), and blocked in PBT + 5% BSA.

Antibodies used were: mouse anti-Wg (DSHB 1:50), mouse anti-Repo (DSHB 1:50), mouse anti-Fz1 (DSHB 1:50), mouse anti-Cyt-Arm (DSHB 1:50), mouse anti-MMP1 (DSHB 5H7B11, 3A6B4, 3B8D12, 1:50), rabbit anti-MMP2 (1:500, K. Broadie) [85], guinea pig anti-grnd (1:250, P. Leopold) [75], mouse anti-β-galactosidase (Sigma, 1:500), rabbit anti-GFP (Invitrogen A11122, 1:500), mouse anti-GFP (Invitrogen A11120, 1:500), mouse anti-brp (DSHB Nc82, 1:20), Rabbit anti-Hrp (Jackson Immunoresearch 111-035-144, 1:400).

Secondary antibodies: anti-mouse Alexa 488, 568, 647, anti-rabbit Alexa 488, 568, 647 (Thermofisher, 1:500). DNA was stained with 2-(4-amidinophenyl)-1H-indole-6-carboxamidine (DAPI, 1μM).

### Western blots

For western blots, we used NuPAGE Bis-Tris Gels 4–12% (Invitrogen), and the following primary antibodies: mouse anti-MMP1 (DSHB 1:500) and mouse anti-tubulin (1:10,000 Sigma), we use Tubulin as a loading control instead of actin because the tumor microtubes are Actin positive and tubulin negative as previously described [10]. There were 3 biological replicates and Relative MMP1 Average pixel intensity was measured using measurement tool from Image Studio Lite Ver 5.2 and normalized against Tubulin.

### Survival assay

Males *Tub-Gal80; Repo-Gal4* were crossed with females bearing a control construct (*UAS–LacZ*) or glioma (*UAS–PI3K*^*dp110*^*; UAS-EGFR*^*λ*^) or glioma + *Mmp1-RNAi* and glioma + *Mmp2-RNAi* and raised at 17°C. Progeny bearing a glioma or glioma + Mmp1-RNAi and glioma + Mmp2-RNAi (experimental) or LacZ (control) chromosomes were put at 29ºC and viability was calculated as the percentage of surviving flies with respect to the starting number of flies as follows: viability = observed (nº of flies)/starting nº of flies × 100. Six independent vials for experimental (*n*= 6) and controls (*n*= 6) were analyzed, with each vial with 10 flies.

### Imaging

Fluorescent labeled samples were mounted in Vectashield mounting media with DAPI (Vector Laboratories) and analyzed by Confocal microscopy (LEICA TCS SP5). Images were processed using Leica LAS AF Lite and Fiji (Image J 1.50e). Images were assembled using Adobe Photoshop CS5.1.

### Quantifications

Relative Wg, Fz1, Cyt-Arm, TRE-RFP, grnd, MMP1 and MMP2 staining within brains was determined from images taken at the same confocal settings. Average pixel intensity was measured using measurement log tool from Fiji 1.51g and Adobe Photoshop CS5.1. Average pixel intensity was measured in the glial tissue and in the adjacent neuronal tissue (N<10 for each sample in triplicates) and expressed as a Glia/Neuron ratio in all cases but Cyt-Arm that was expressed as Neuron/Glia ratio. Glial network volume was quantified using Imaris surface tool (Imaris 6.3.1 software).

For the co-localization of egr-GFP in glial cells, GFP channel volume was quantified using Imaris surface tool. We selected a specific threshold for the total volume in the control samples and then we applied these conditions to the analysis of the corresponding experimental sample. Then we applied a co-localization filter (intensity center of the red channel)

The number of Repo^+^ cells, the number of synaptic active sites and the number of puc-lacZ positive cells was quantified by using the spots tool Imaris 6.3.1 software, we selected a minimum size and threshold for the puncta in the control samples of each experiment. Then we applied these conditions to the analysis of each corresponding experimental sample. For the puc-lacZ glia or neuron co-localization studies we quantified the total number of puc-lacZ^+^ cells and then applied a co-localization filter (intensity center of the channel of interest) using the Spots tool from the Imaris 6.3.1 Bitplane Scientific Solutions software.

### Statistical Analysis

To analyze and plot the data, we used Microsoft Excel 2013 and GraphPad Prism 6. We performed a D’Agostino & Pearson normality test and the data found to have a normal distribution were analyzed by a two-tailed t test with Welch-correction. In the case of multiple comparisons, we used a One-way ANOVA with Bonferroni post-test. The data that did not pass the normality test were subjected to a two-tailed Mann–Whitney U test or in the case of multiple comparisons a Kruskal–Wallis test with Dunns post-test. Error bars represent standard error of the mean. * represents p value ≤ .05; ** p value ≤ .01; *** p value ≤ .001. Statistical values of p value > .05 were not considered significant, (n.s.).

