## Supplemental Figure legends for "Wingless promotes JNK/MMPs positive feedback loop mediate tumour microtubes expansion, glioma progression and neurodegeneration"

**Figure S1: MMP2 is upregulated in GB**

Brains from 3rd instar larvae displayed at the same scale. Glia is labelled with *UAS-Ihog-RFP* driven by *repo-Gal4* to visualize active cytonemes/ TM structures in glial cells, and stained with MMP2 (green). (A) MMP2 is homogeneously distributed in control sections, with a slight accumulation in the Ihog+ projections (B) MMP2 accumulates in the TMs and specifically in the projections that are in contact with the neuronal clusters. (C) Inhibition of *Gap43* by *RNAi* in glioma brains restores a normal glial network and MMP1 does not accumulate showing a homogeneous staining along the brain section. (D) Inhibition of *Fz1* by *RNAi* in glioma brains restores a normal MMP2 distribution. MMP2 does not accumulate showing a homogeneous staining along the brain section. Nuclei are marked with DAPI. (E) Quantification of MMP2 staining ratio between $ihog^+$ and $ihog^-$ domains. (F) MMP2-GFP reporter (green) showing activation in the glioma cell membranes (red). (G-I) Glial cell bodies and membranes labelled in green (CD2-GFP) driven by *repo-Gal4* to the glial cells and stained with MMP2 (red). (G) MMP2 is homogeneously distributed in control sections. (H) MMP2 accumulates in the glial cells upon Fz1 overexpression. (I) Quantification of MMP2 staining ratio between $GFP^+$ and $GFP^-$ domains. Nuclei are marked with DAPI. Error bars show S.D. * P<0.01, ** P<0.001, *** P<0.0001 or ns for non-significant. Scale bar size is indicated in this and all figures.

**Figure S2: Independent constitutive activation of PI3K or EGFR or ectopic *dmyc* are not responsible for MMP1 accumulation.**

Brains from 3rd instar larvae. Glia is labelled with *UAS-Ihog-RFP* driven by *repo-Gal4* to visualize active cytonemes/ TM structures in glial cells, and stained with MMP1 (green). (A-C) (A) MMP1 (green) is homogeneously distributed in $dp110^{CAAX}$ sections and the glioma network shown in red by Ihog-RFP does not overgrow or encapsulate neuronal clusters. (B) Fz1 (green) is homogeneously distributed in $TOR\text{-}DER^{CA}$ sections and the glioma network shown in red by Ihog-RFP does not overgrow or encapsulate neuronal clusters, similar to dmyc overexpression in glial cells (C). Nuclei are marked with DAPI. (D) Quantification of MMP1 staining ratio between $ihog^+$ and $ihog^-$ domains. Error bars show S.D. * P<0.01, ** P<0.001, *** P<0.0001 or ns for non-significant. Scale bar size are indicated in this and all figures.

**Figure S3: Egr and its receptor Grindelwald accumulate in GB**

Brains from 3rd instar larvae. Glia is labelled with *UAS-Ihog-RFP* driven by *repo-Gal4* to visualize active cytonemes/ TM structures in glial cells, carry a egr-GFP reporter (green). (A-B) In control brain sections egr-GFP (green) signal is localized mostly (65%) in neurons in close contact with glial cells. (C-D) In glioma brain sections egr-GFP (green) signal shifts and is localized mostly (55%) in glioma cells. (E) Quantification of the percentage of Egr in glia and neurons in control and glioma samples. (F-G) grnd staining (green) in control and glioma samples. (H) Quantification of the grnd pixel intensity. Nuclei are marked with DAPI. Error bars show S.D. * P<0.01, ** P<0.001, *** P<0.0001 or ns for non-significant. Scale bar size is indicated in this and all figures.

**Figure S4: JNK mediates GB progression**

Brains from 3rd instar larvae displayed at the same scale. Glia are labeled with *UAS-Ihog-RFP* driven by *repo-Gal4* to visualize active cytonemes/ TM structures in glial cells, and stained with Repo (green) in the following genotypes (A-G) control, $bsk^{DN}$, glioma $bsk^{DN}$, $egr^{-/-}$ and Glioma $egr^{-/-}$, $grnd^{EXTRA}$, glioma $grnd^{EXTRA}$ brain sections. The number of $Repo^+$ cells is quantified in (H). Nuclei are marked with DAPI. Error bars show S.D. * P<0.01, ** P<0.001, *** P<0.0001 or ns for non-significant. Scale bar size are indicated in this and all figures.

**Figure S5: JNK mediates Fz1 accumulation in TMs**

Brains from 3rd instar larvae displayed at the same scale. Glia are labeled with *UAS-Ihog-RFP* driven by *repo-Gal4* to visualize active cytonemes/ TM structures in glial cells, and stained with Fz1 (green) in the following genotypes (A-E) control, glioma, glioma Gap43-RNAi, Glioma Fz1-RNAi and glioma $bsk^{DN}$ brain sections. Fz1 average pixel intensity quantification ratio between $ihog^{+}$ and $ihog^{-}$ domains (F). Nuclei are marked with DAPI. Error bars show S.D. * P<0.01, ** P<0.001, *** P<0.0001 or ns for non-significant. Scale bar size is indicated in this and all figures.

**Figure S6: Validation of tools, RNAis and antibodies**

(A) *UAS-MMP1-RNAi* tool was validated in epithelial tissue, wing imaginal discs. Upon *UAS-MMP1-RNAi* expression in the posterior compartment (marked with GFP), there is a reduction in MMP1 protein signal compared with the anterior compartment from the same tissue. (B) *UAS-MMP2-RNAi* tool was validated in epithelial tissue, wing imaginal discs. Upon *UAS-MMP2-RNAi* expression in the posterior compartment (marked with GFP), there is a reduction in MMP2 protein signal compared with the anterior compartment from the same tissue.

**Figure S7: Wg accumulation in TMs, activation of wg signaling pathway and glial cell number increase in GB requires MMPs**

Brains from 3rd instar larvae displayed at the same scale. Glia are labeled with *UAS-Ihog-RFP* driven by *repo-Gal4* to visualize active cytonemes/ TM structures in glial cells, and stained with Wg (A-F) or Cyt-Arm (H-M) in green in the following genotypes control, glioma, MMP1-RNAi, glioma MMP1-RNAi, MMP2-RNAi and glioma MMP2-RNAi brain sections. (G) Quantification of Wg average pixel intensity staining ratio between $ihog^{+}$ and $ihog^{-}$ domains. (N) Quantification of Cyt-Arm average pixel intensity Neuron/Glia ratio between $ihog^{-}$ and $ihog^{+}$ domains. Nuclei are marked with DAPI. Error bars show S.D. * P<0.01, ** P<0.001, *** P<0.0001 or ns for non-significant. Scale bar size is indicated in this and all figures.

**Genotypes**

**Figure 1**

(A) *repo-Gal4, ihog-RFP/UAS-lacZ*

(B) *UAS-dEGFR*$^{\lambda}$*, UAS-dp110*$^{CAAX}$*;; repo-Gal4, UAS-ihog-RFP*

(C) *UAS-dEGFR*$^{\lambda}$*, UAS-dp110*$^{CAAX}$*;; repo-Gal4, UAS-ihog-RFP /UAS-Gap43-RNAi*

(D) *UAS-dEGFR*$^{\lambda}$*, UAS-dp110*$^{CAAX}$*; /UAS-Fz1-RNAi; repo-Gal4, UAS-ihog-RFP*

**Figure 2**

(A-B) *w;; Repo-LexA, LexAop-CD2-GFP/CyO*

(C-D) *w;; Repo-LexA, LexAop-CD2-GFP/LexAop-Fz1*

**Figure 3**

(A) *Gal80*$^{ts}$*/repo-Gal4, myr-RFP/puc-lacZ*

(B-C) *Gal80*$^{ts}$*/UAS-dEGFR*$^{\lambda}$*, UAS-dp110*$^{CAAX}$*;; repo-Gal4, myr-RFP/puc-lacZ*

(E) *TRE-RFP; repo-Gal4, UAS-GFP*$^{NLS}$*/UAS-lacZ*

(F) *UAS-dEGFR*$^{\lambda}$*, UAS-dp110*$^{CAAX}$*; TRE-RFP; repo-Gal4, UAS-GFP*$^{NLS}$

(G) *UAS-dEGFR*$^{\lambda}$*, UAS-dp110*$^{CAAX}$*; TRE-RFP; repo-Gal4, UAS-GFP*$^{NLS}$*/UAS-Gap43-RNAi*

(H) *UAS-dEGFR*$^{\lambda}$*, UAS-dp110*$^{CAAX}$*; TRE-RFP/UAS-Fz1-RNAi; repo-Gal4, UAS-GFP*$^{NLS}$

**Figure 4**

(A) *repo-Gal4, ihog-RFP/UAS-lacZ*

(B) *UAS-dEGFR*$^{\lambda}$*, UAS-dp110*$^{CAAX}$*;; repo-Gal4, UAS-ihog-RFP*

(C) *UAS-grnd-RNAi; repo-Gal4, ihog-RFP*

(D) *UAS-dEGFR*$^{\lambda}$*, UAS-dp110*$^{CAAX}$*; UAS-grnd-RNAi; repo-Gal4, UAS-ihog-RFP*

(E) *grnd*$^{MINOS}$*/grnd*$^{MINOS}$*; repo-Gal4, ihog-RFP*

(F) *UAS-dEGFR*$^{\lambda}$*, UAS-dp110*$^{CAAX}$*; grnd*$^{MINOS}$*/grnd*$^{MINOS}$*; repo-Gal4, UAS-ihog-RFP*

**Figure 5**

(A) *repo-Gal4, ihog-RFP/UAS-lacZ*

(B) *UAS-dEGFR*$^{\lambda}$*, UAS-dp110*$^{CAAX}$*;; repo-Gal4, UAS-ihog-RFP*

(C) *UAS-grnd-RNAi; repo-Gal4, ihog-RFP*

(D) *UAS-dEGFR*$^{\lambda}$*, UAS-dp110*$^{CAAX}$*; UAS-grnd-RNAi; repo-Gal4, UAS-ihog-RFP*

(E) *grnd*$^{MINOS}$*/grnd*$^{MINOS}$*; repo-Gal4, ihog-RFP*

(F) *UAS-dEGFR*$^{\lambda}$*, UAS-dp110*$^{CAAX}$*; grnd*$^{MINOS}$*/grnd*$^{MINOS}$*; repo-Gal4, UAS-ihog-RFP*

**Figure 6**

(A) *repo-Gal4, ihog-RFP/UAS-lacZ*

(B) *UAS-dEGFR$^{\lambda}$, UAS-dp110$^{CAAX}$;; repo-Gal4, UAS-ihog-RFP*

(C) *UAS-grnd-RNAi; repo-Gal4, ihog-RFP*

(D) *UAS-dEGFR$^{\lambda}$, UAS-dp110$^{CAAX}$; UAS-grnd-RNAi; repo-Gal4, UAS-ihog-RFP*

(E) *grnd$^{MINOS}$/grnd$^{MINOS}$; repo-Gal4, ihog-RFP*

(F) *UAS-dEGFR$^{\lambda}$, UAS-dp110$^{CAAX}$; grnd$^{MINOS}$/grnd$^{MINOS}$; repo-Gal4, UAS-ihog-RFP*

**Figure 7**

(A) *repo-Gal4, ihog-RFP/UAS-lacZ*

(B) *UAS-dEGFR$^{\lambda}$, UAS-dp110$^{CAAX}$;; repo-Gal4, UAS-ihog-RFP*

(C) *UAS-dEGFR$^{\lambda}$, UAS-dp110$^{CAAX}$;; repo-Gal4, UAS-ihog-RFP/ UAS-bsk$^{DN}$*

(D) *UAS-grnd-RNAi; repo-Gal4, ihog-RFP*

(E) *UAS-dEGFR$^{\lambda}$, UAS-dp110$^{CAAX}$; UAS-grnd-RNAi; repo-Gal4, UAS-ihog-RFP*

**Figure 8**

(A) *repo-Gal4, ihog-RFP/UAS-lacZ*

(B) *UAS-dEGFR$^{\lambda}$, UAS-dp110$^{CAAX}$;; repo-Gal4, UAS-ihog-RFP*

(C) *UAS-MMP1-RNAi; repo-Gal4, ihog-RFP*

(D) *UAS-dEGFR$^{\lambda}$, UAS-dp110$^{CAAX}$; UAS-MMP1-RNAi; repo-Gal4, UAS-ihog-RFP*

(E) *UAS-MMP2-RNAi; repo-Gal4, ihog-RFP*

(F) *UAS-dEGFR$^{\lambda}$, UAS-dp110$^{CAAX}$; UAS-MMP2-RNAi; repo-Gal4, UAS-ihog-RFP*

**Figure 9**

(A) *repo-Gal4, ihog-RFP/UAS-lacZ*

(B) *UAS-dEGFR$^{\lambda}$, UAS-dp110$^{CAAX}$;; repo-Gal4, UAS-ihog-RFP*

(C) *UAS-MMP1-RNAi; repo-Gal4, ihog-RFP*

(D) *UAS-dEGFR$^{\lambda}$, UAS-dp110$^{CAAX}$; UAS-MMP1-RNAi; repo-Gal4, UAS-ihog-RFP*

(E) *UAS-MMP2-RNAi; repo-Gal4, ihog-RFP*

(F) *UAS-dEGFR$^{\lambda}$, UAS-dp110$^{CAAX}$; UAS-MMP2-RNAi; repo-Gal4, UAS-ihog-RFP*

**Figure S1**

(A) *repo-Gal4, ihog-RFP/UAS-lacZ*

(B) *UAS-dEGFR$^{\lambda}$, UAS-dp110$^{CAAX}$;; repo-Gal4, UAS-ihog-RFP*

(C) *UAS-dEGFR$^{\lambda}$, UAS-dp110$^{CAAX}$;; repo-Gal4, UAS-ihog-RFP /UAS-Gap43-RNAi*

(D) *UAS-dEGFR$^{\lambda}$, UAS-dp110$^{CAAX}$; /UAS-Fz1-RNAi; repo-Gal4, UAS-ihog-RFP*

(F) *Gal80$^{ts}$/UAS-dEGFR$^{\lambda}$, UAS-dp110$^{CAAX}$; MMP2-GFP; repo-Gal4, myr-RFP*

(G) *w;; Repo-LexA, LexAop-CD2-GFP/CyO*

(H) *w;; Repo-LexA, LexAop-CD2-GFP/LexAop-Fz1*

**Figure S2**

(A) *UAS-dp110$^{CAAX}$;; repo-Gal4, UAS-ihog-RFP*

(B) *UAS-TOR-DER$^{CA}$ ;; repo-Gal4, UAS-ihog-RFP*

(C) *repo-Gal4, UAS-ihog-RFP/UAS-dmyc*

**Figure S3**

(A-B) egr-GFP; *repo-Gal4, ihog-RFP/UAS-lacZ*

(C-D) *UAS-dEGFR$^{\lambda}$, UAS-dp110$^{CAAX}$;egr-GFP; repo-Gal4, UAS-ihog-RFP*

(F) *repo-Gal4, ihog-RFP/UAS-lacZ*

(G) *UAS-dEGFR$^{\lambda}$, UAS-dp110$^{CAAX}$;; repo-Gal4, UAS-ihog-RFP*

**Figure S4**

(A) *repo-Gal4, ihog-RFP/UAS-lacZ*

(B) *repo-Gal4, ihog-RFP/ UAS-bsk$^{DN}$*

(C) *UAS-dEGFR$^{\lambda}$, UAS-dp110$^{CAAX}$;; repo-Gal4, UAS-ihog-RFP/ UAS-bsk$^{DN}$*

(D) *egr$^{-}$/egr$^{-}$ ; repo-Gal4, ihog-RFP*

(E) *UAS-dEGFR$^{\lambda}$, UAS-dp110$^{CAAX}$; egr$^{-}$/egr$^{-}$; repo-Gal4, UAS-ihog-RFP*

(F) *UAS-grnd$^{EXTRA}$/ repo-Gal4, ihog-RFP*

(G) *UAS-dEGFR$^{\lambda}$, UAS-dp110$^{CAAX}$; UAS-grnd$^{EXTRA}$/repo-Gal4, UAS-ihog-RFP*

**Figure S5**

(A) *repo-Gal4, ihog-RFP/UAS-lacZ*

(B) *UAS-dEGFR$^{\lambda}$, UAS-dp110$^{CAAX}$;; repo-Gal4, UAS-ihog-RFP*

(C) *UAS-dEGFR$^{\lambda}$, UAS-dp110$^{CAAX}$;; repo-Gal4, UAS-ihog-RFP /UAS-Gap43-RNAi*

(D) *UAS-dEGFR$^{\lambda}$, UAS-dp110$^{CAAX}$; /UAS-Fz1-RNAi; repo-Gal4, UAS-ihog-RFP*

(E) *UAS-dEGFR$^{\lambda}$, UAS-dp110$^{CAAX}$;; repo-Gal4, UAS-ihog-RFP/ UAS-bsk$^{DN}$*

**Figure S6**

(A) *Gal80$^{ts}$/UAS-MMP1-RNAi; en-Gal4, UAS-GFP*

(B) *Gal80$^{ts}$/UAS-MMP2-RNAi; en-Gal4, UAS-GFP*

**Figure S7**

(A, H) *repo-Gal4, ihog-RFP/UAS-lacZ*

(B, I) *UAS-dEGFR$^{\lambda}$, UAS-dp110$^{CAAX}$;; repo-Gal4, UAS-ihog-RFP*

(C, J) *UAS-MMP1-RNAi; repo-Gal4, ihog-RFP*

(D, K) *UAS-dEGFR$^{\lambda}$, UAS-dp110$^{CAAX}$; UAS-MMP1-RNAi; repo-Gal4, UAS-ihog-RFP*

(E, L) *UAS-MMP2-RNAi; repo-Gal4, ihog-RFP*

(F, M) *UAS-dEGFR*$^{\lambda}$*, UAS-dp110*$^{CAAX}$*; UAS-MMP2-RNAi; repo-Gal4, UAS-ihog-RFP*
