## Supplementary figures and images for "Wingless promotes JNK/MMPs positive feedback loop mediate tumour microtubes expansion, glioma progression and neurodegeneration"

### Figure S1

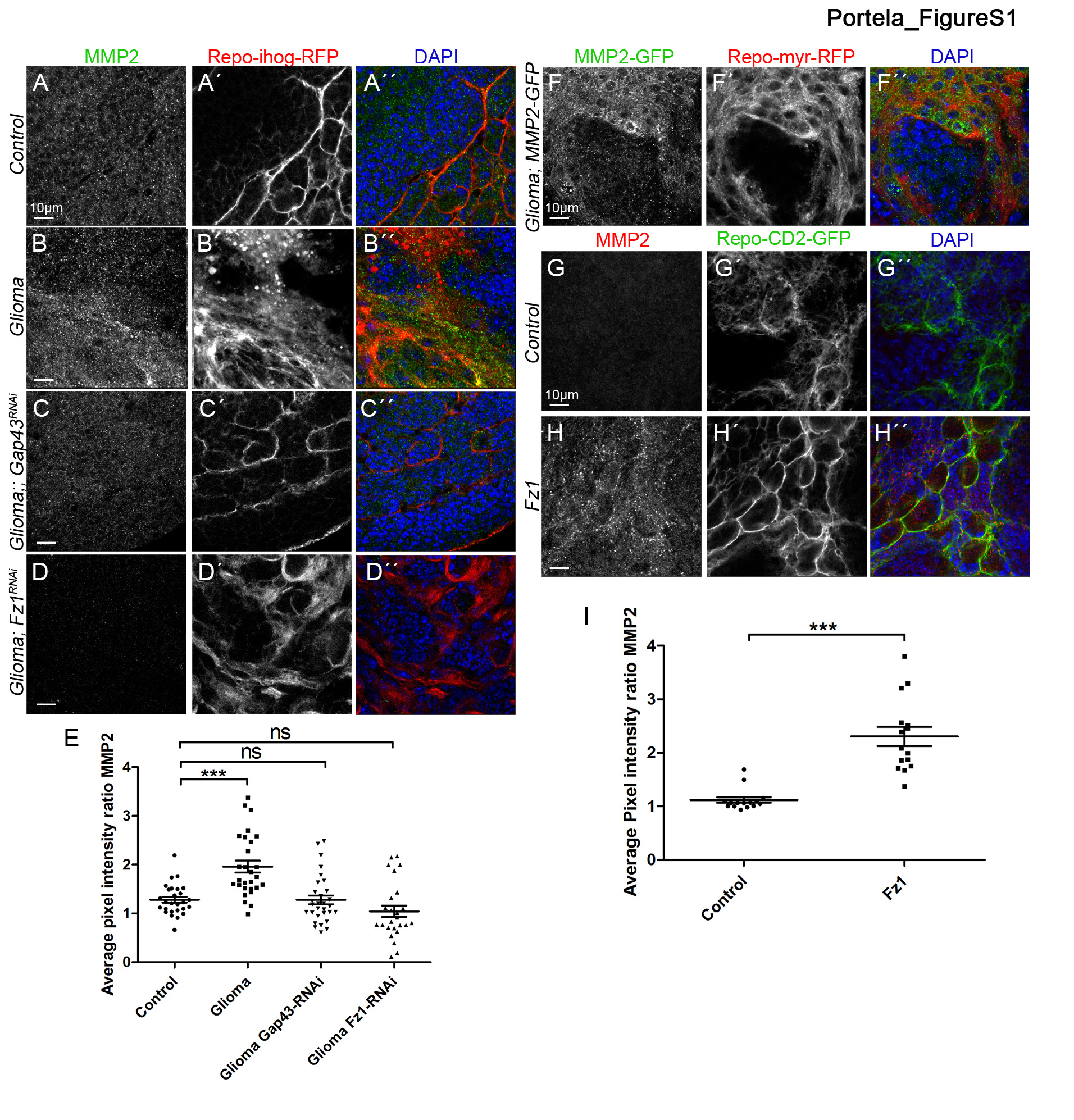

### Figure S2

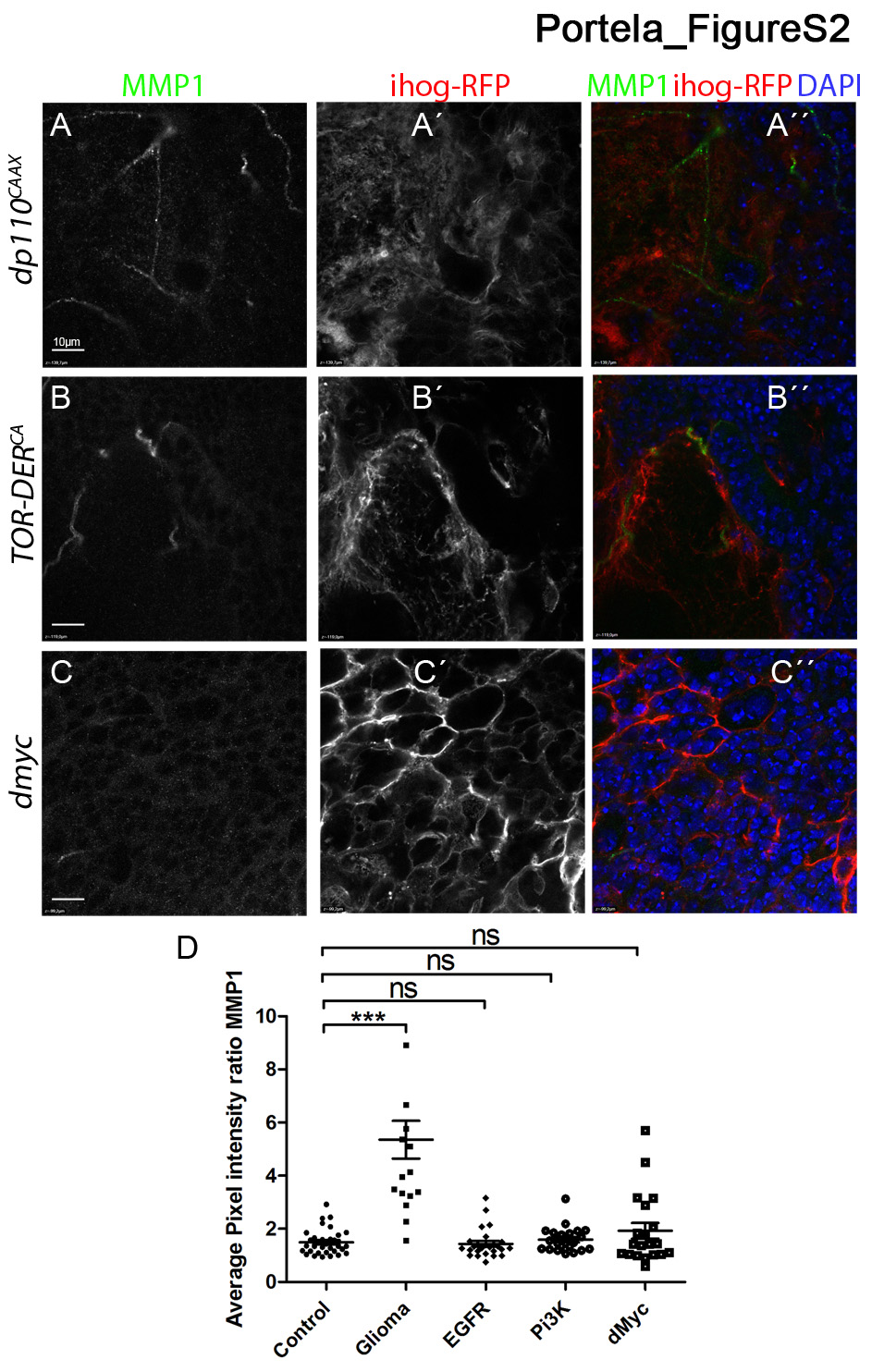

### Figure S3

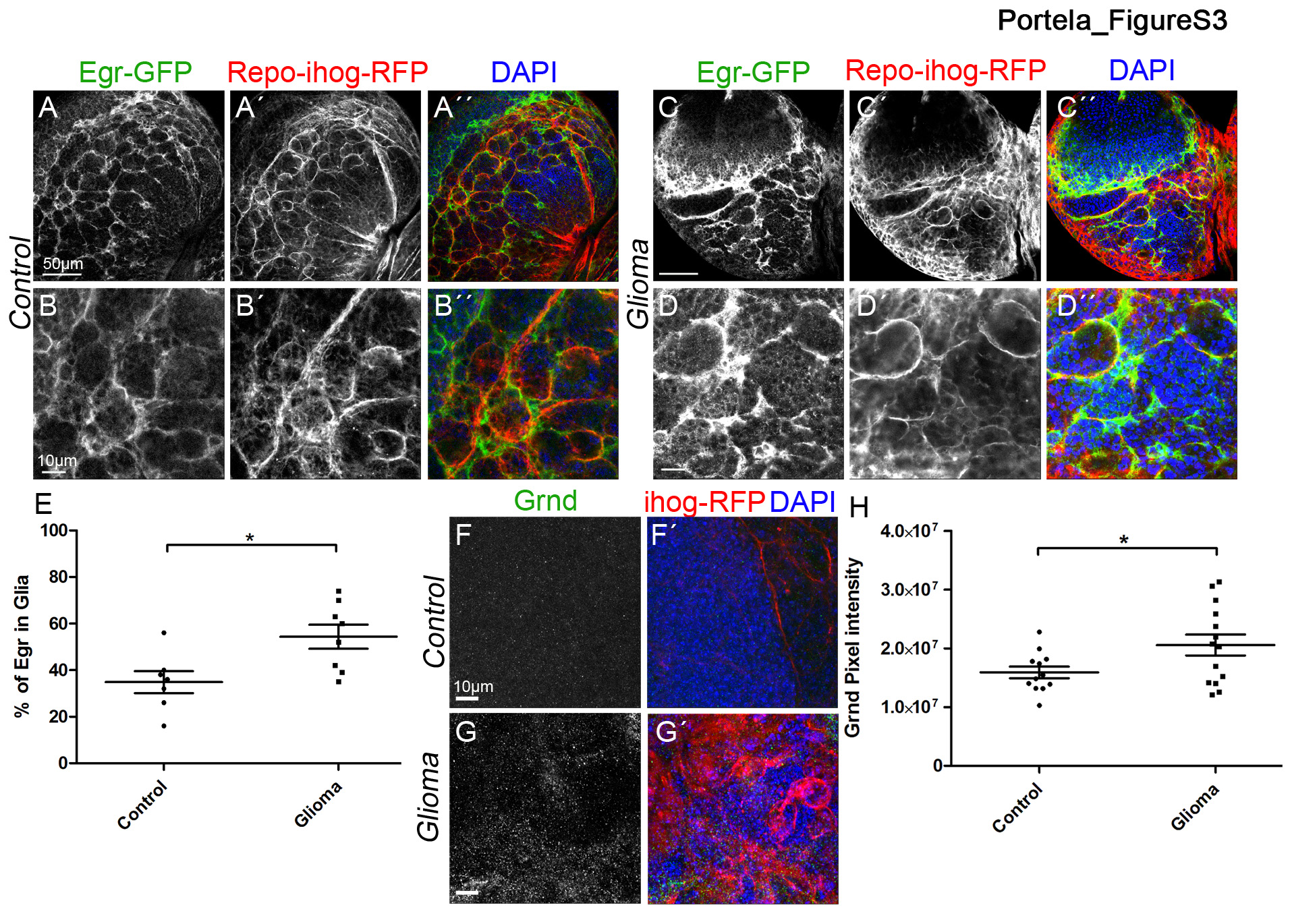

### Figure S4

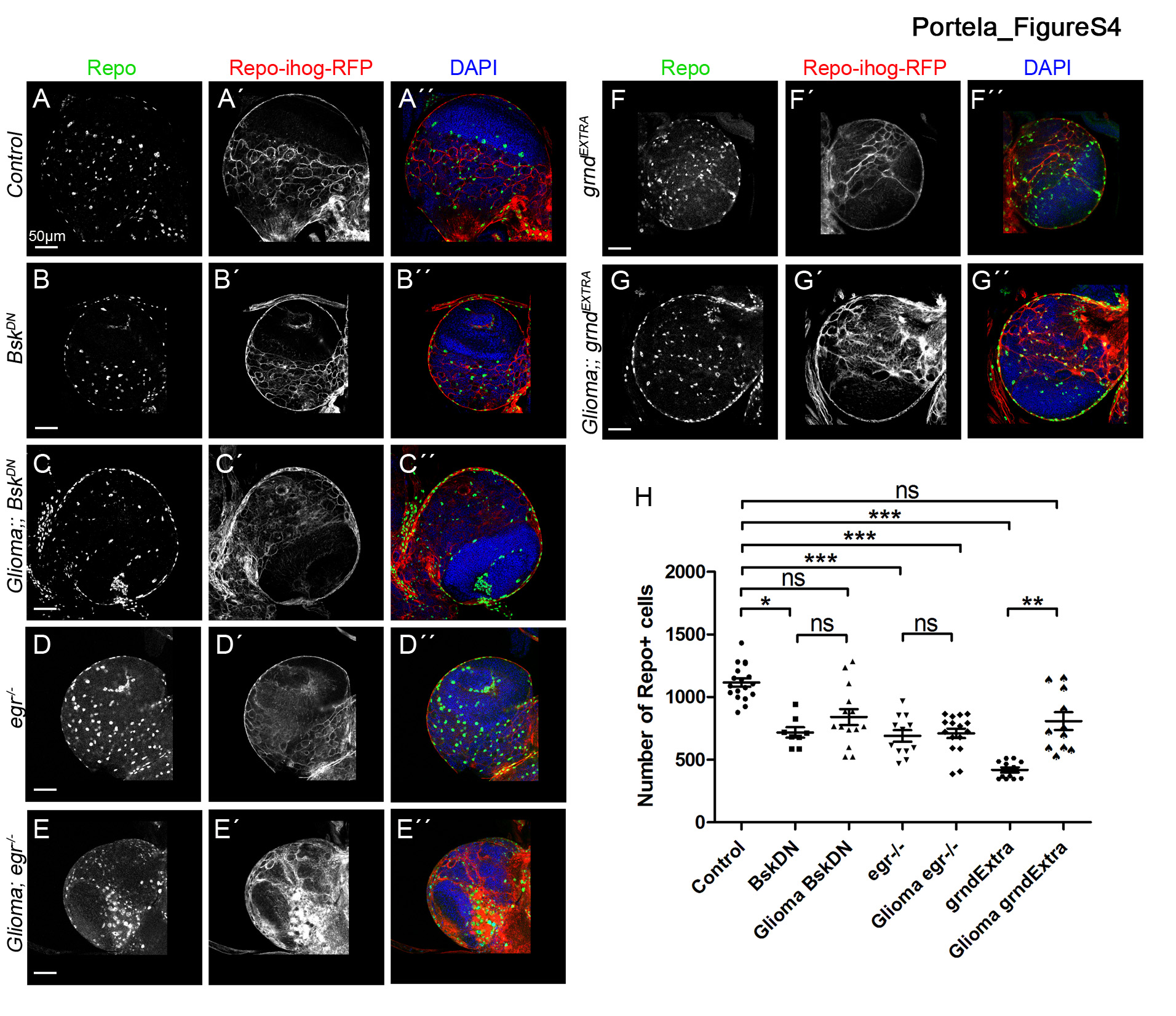

### Figure S5

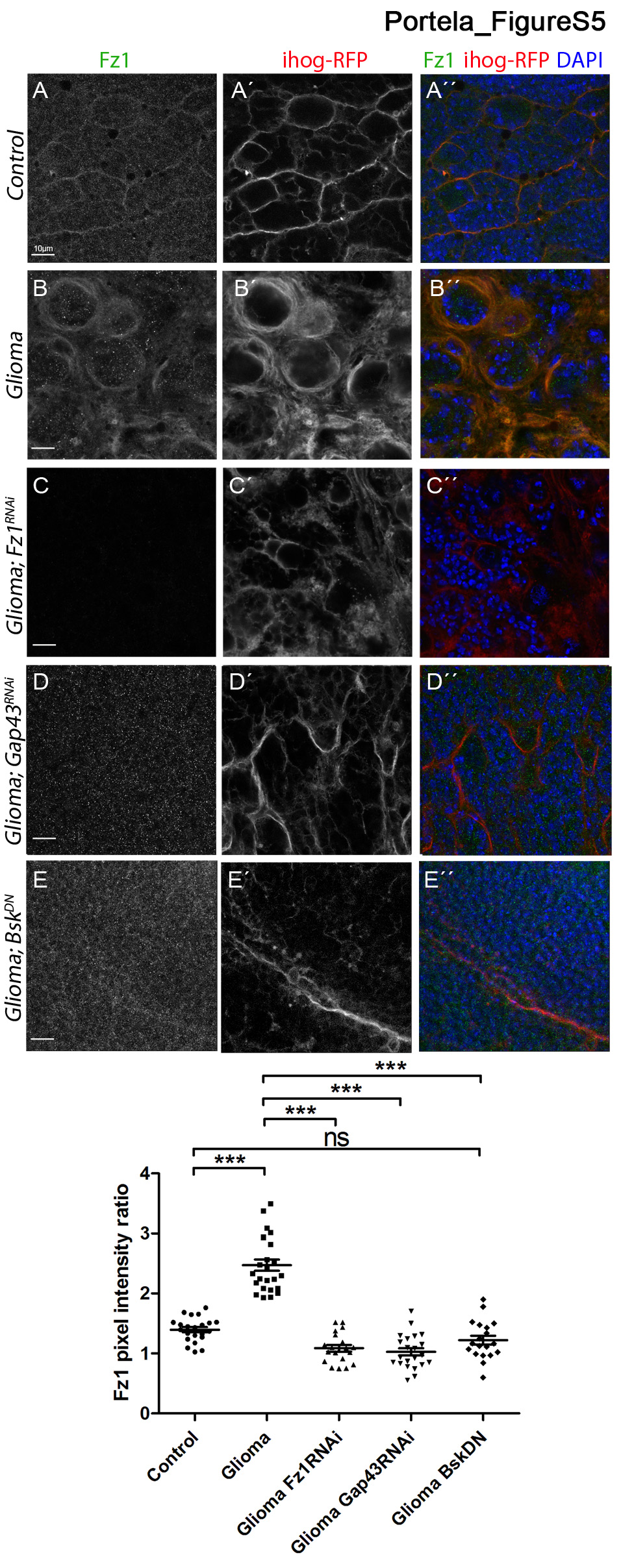

### Figure S6

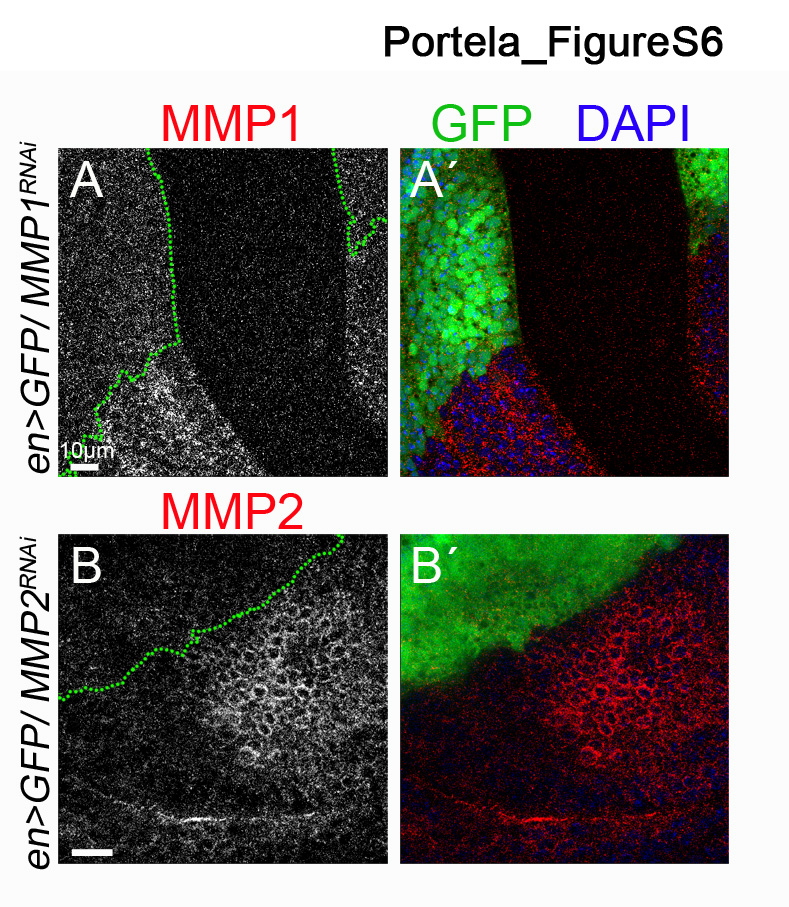

### Figure S7

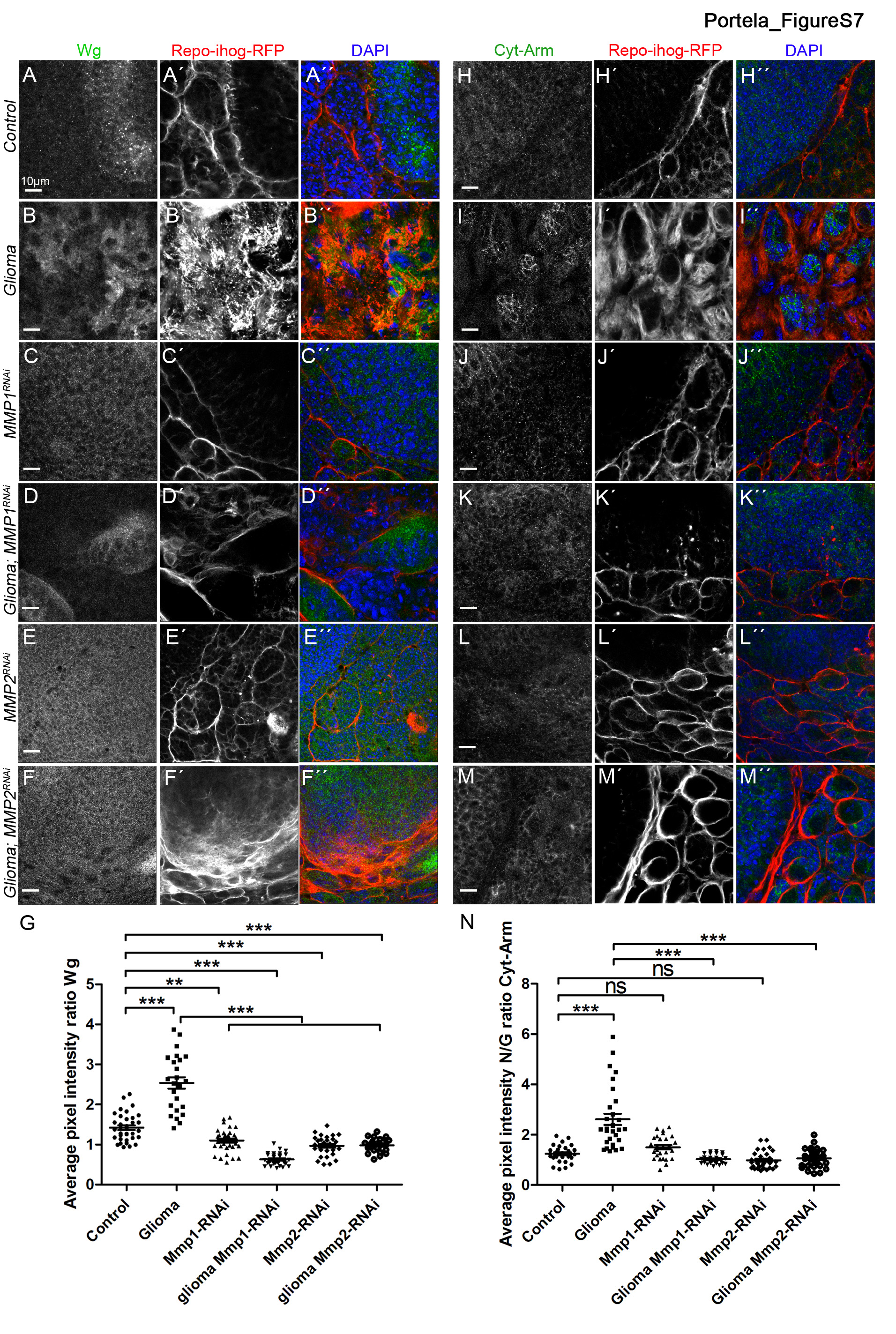
